# Nanoengineered DNA origami with repurposed TOP1 inhibitors targeting myeloid cells for the mitigation of neuroinflammation

**DOI:** 10.1101/2021.10.04.462880

**Authors:** Keying Zhu, Yang Wang, Heela Sarlus, Keyi Geng, Erik Nutma, Jingxian Sun, Shin-Yu Kung, Cindy Bay, Jinming Han, Harald Lund, Sandra Amor, Jun Wang, Xingmei Zhang, Claudia Kutter, André Ortlieb Guerreiro Cacais, Björn Högberg, Robert A. Harris

**Affiliations:** Applied Immunology and Immunotherapy, Department of Clinical Neuroscience, Karolinska Institutet, Center for Molecular Medicine, Karolinska University Hospital, 171 76 Stockholm, Sweden; Department of Medical Biochemistry and Biophysics, Karolinska Institutet, 171 77 Stockholm, Sweden; Department of Microbiology, Tumor and Cell Biology, Science for Life Laboratory, Karolinska Institute, 171 77 Stockholm, Sweden; Department of Pathology, Amsterdam UMC, Location VUmc, The Netherlands; Department of Integrative Medicine and Neurobiology, School of Basic Medical Sciences; State Key Laboratory of Medical Neurobiology and MOE Frontiers Center for Brain Science; Shanghai Medical College, Fudan University, Shanghai, 200032, China; Department of Physiology and Pharmacology, Karolinska Institutet, Center for Molecular Medicine, Karolinska University Hospital, 171 76 Stockholm, Sweden

## Abstract

Targeting myeloid cells, especially microglia, for the treatment of neuroinflammatory diseases such as multiple sclerosis (MS), is underappreciated. Here, we screened a library of compounds and identified the topoisomerase 1 (TOP1) inhibitor camptothecin (CPT) as a promising drug candidate for microglial modulation. CPT and its FDA-approved analog topotecan (TPT) inhibited inflammatory responses in microglia and macrophages, and ameliorated neuroinflammation in mice. Transcriptomic analysis of sorted microglia revealed an altered transcriptional phenotype following TPT treatment, with Ikzf1 identified as a potential target. Importantly, TOP1 expression was found elevated in several neuroinflammatory conditions, including human MS brains. To achieve targeted delivery to myeloid cells we designed a nanosystem using DNA origami and loaded TPT into it (TopoGami). TopoGami also significantly suppressed the inflammatory response in microglia and mitigated disease progression in MS-like mice. Our findings suggest that TOP1 inhibition represents a therapeutic strategy for neuroinflammatory diseases, and the proposed nanosystem may foster future research and drug development with a demand to target myeloid cells.

## Introduction

The functions of the innate immune system are predominantly executed by cells of the myeloid lineage (‘myeloid cells’), which comprise monocytes, dendritic cells, macrophages, and microglia. Myeloid cells play important roles in the regulation of inflammation, non-specific host defense against microbes, the clearance of tissue debris, and the remodeling of tissues.

Neuroinflammation refers to inflammatory responses occurring within the central nervous system (CNS) and is a common feature of various disease states or etiology that may be acute or chronic. During CNS neuroinflammatory conditions such as multiple sclerosis (MS), amyotrophic lateral sclerosis (ALS), and rodent experimental autoimmune encephalomyelitis (EAE), a variety of myeloid cell subtypes infiltrate the CNS, thereby disturbing the homeostasis.

Emerging evidence has indicated a pivotal role of myeloid cells in the immunopathology of neurological diseases. Infiltrating monocyte-derived macrophages destroy neurons and oligodendrocytes during MS, and the action of pro-inflammatory microglia/macrophages drives motor neuron death in ALS. A recent human genome-wide association study including 47,429 MS patients and 68,374 control cases determined that MS susceptibility genes were enriched in microglia, but not in astrocytes or neurons, suggesting a potential but overlooked role of microglia in MS susceptibility (*1*). The *C9orf72* gene, whose dysregulation accounts for the most frequent genetic cause of familial ALS, is highly expressed in myeloid cells (*2*). The absence of *C9orf72* in myeloid cells disturbs homeostasis and predisposes individuals to neuroinflammatory disorders (*3*). A recent study also demonstrated that although peripheral myeloid cell infiltration to the CNS was limited, macrophages along peripheral axons of motor neurons are activated in ALS mouse models and ALS patients. Replacing them with less inflammatory macrophages reduced the activation of microglia, delayed symptoms, and increased survival in ALS mice (*4*). Myeloid cells therefore clearly represent a crucial therapeutic target for neuroinflammatory diseases.

While endowed with high translational value and clinical demand, therapeutics developed to specifically target myeloid cells are scarce. In this study, we designed a drug screening strategy using *Connectivity Map* (CMAP) to identify drugs with the potential to modulate microglial function. The topoisomerase 1 (TOP1) inhibitor camptothecin (CPT) emerged as a potent agent to regulate microglial activation. We confirmed the beneficial effect of CPT and its FDA-approved analog topotecan (TPT) on neuroinflammatory conditions and determined an increased TOP1 expression during neuroinflammation. Lastly, to achieve enhanced, specific delivery of TPT to myeloid cells and to circumvent potential off-target effects on other immune cells, we loaded TPT into a wireframe DNA origami structured in a hexagonal prism (*5, 6*), and surface-coated it with β-glucan (herein referred to as TopoGami). The nanoengineered TopoGami significantly reduced neuroinflammation in a similar fashion to TPT. Our findings indicate the FDA-approved drug topotecan could be repurposed and translated for treating neuroinflammation, and the β-glucan-coated DNA origami system will be of great value for future drug development or studies using demanding a specific targeting of myeloid cells.

## Results

### Identification of TOP1 inhibitors as potent microglial modulators

To find suitable compounds that alter microglia states we used a repurposing approach by utilizing a *Connectivity Map* (CMAP) discovery strategy (*7*), thus enabling a relatively quicker translation of an existing approved drug to a new clinical application. We analyzed a dataset that contains transcriptomic data from human microglia stimulated with defined stimulants to induce different phenotypes: M1(LPS/IFNγ), M2a (IL-4/IL-13), M2c (-4/IL-13/IL-10), and Mtgfb (TGF-β) (Fig. 1a). Although the nomenclature of M1/M2 microglial activation is now considered to be an oversimplified dogma overlooking the heterogeneity and complexity of microglial activation states, *in vitro* stimulation of microglia with defined stimuli clearly distinguishes the proinflammatory M1 phenotype from immunoregulatory M2-subphenotypes (M2a/M2c/Mtgfb). We will use the terminology as it relates to that used in the dataset.

**Fig. 1.**
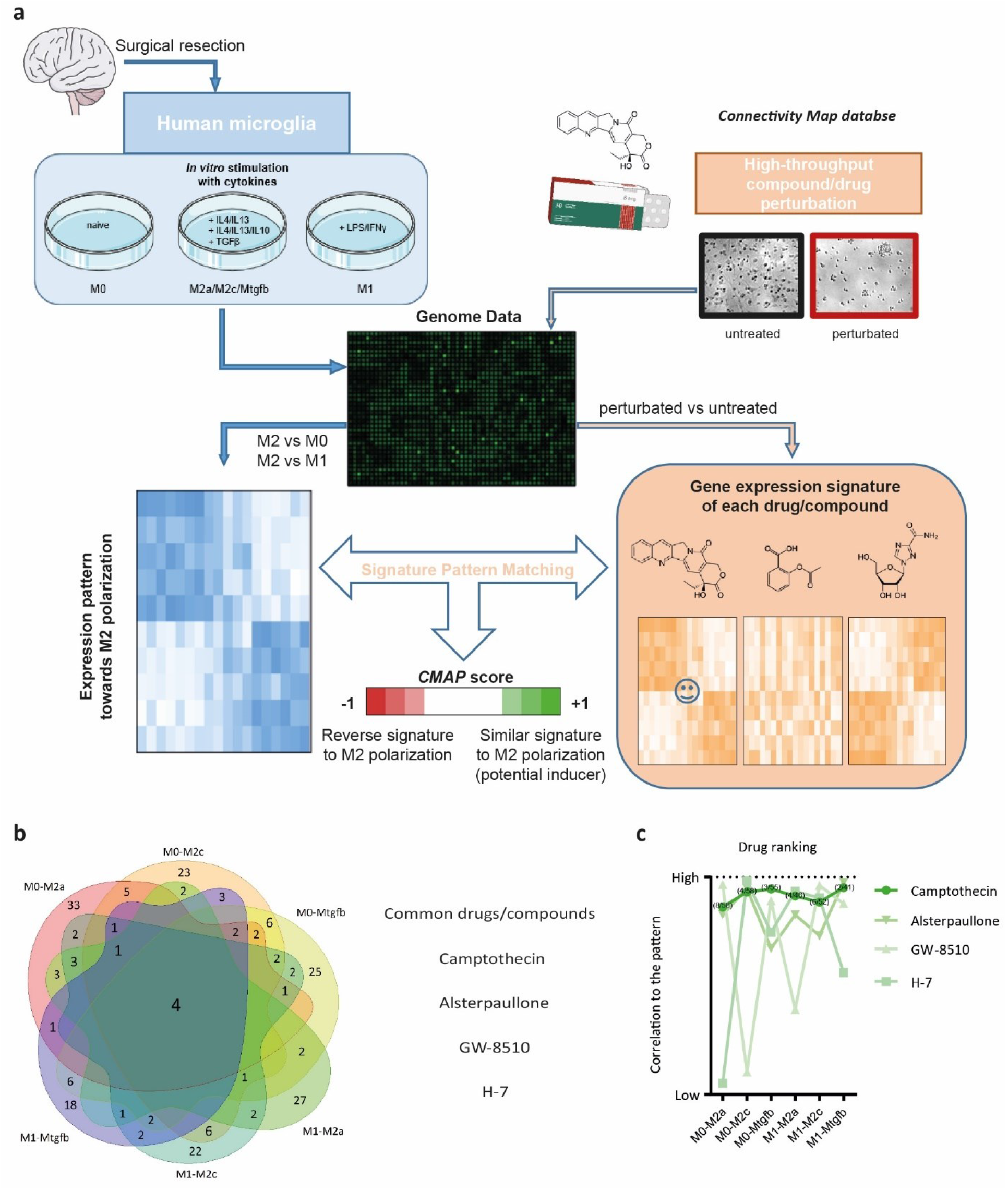
Connectivity Map-based drug screening and repurposing. **a**, A schematic overview of the screening process. A dataset (GSE76737) containing human microglia stimulated with indicated cytokines to induce different phenotypes was analyzed. The differential gene expression profiles from patterns M0 to M2a/M2c/Mtgfb and M1 to M2a/M2c/Mtgfb were identified respectively using the NCBI GEO2R tool. Top 1000 differential genes were selected, and the up-regulated genes and the down-regulated genes were converted to identifiers based on Affymetrix HG-U133A chip, followed by analysis and matching with Connectivity Map. **b**, 4 overlapping compounds/drugs were identified in each of the 6 patterns: camptothecin, alsterpaullone, GW-8510, and H-7. **c**, Compared to the other drug candidates, camptothecin has the highest overall correlation to induce all these 6 patterns.

The six differential gene expression patterns from resting microglia (M0) to M2a/M2c/Mtgfb and M1 to M2a/M2c/Mtgfb activation states were determined. Four compounds (camptothecin, alsterpaullone, GW-8510, and H-7) were identified in the positively correlated drug list in each of these six patterns, indicating they might reprogram microglia into an immunoregulatory phenotype (Fig. 1b). Compared to the other three identified compounds, camptothecin (CPT) had an overall high CMAP score among all the six patterns, and its mechanism of action as a topoisomerase 1 (TOP1) inhibitor is well understood. In addition, derivatives of CPT such as topotecan and irinotecan have been FDA approved for clinical use. We have also listed other potential compounds and drugs from the CMAP screening that might skew microglia towards an immunoregulatory phenotype (Table 1). Surprisingly, a very recent study has also highlighted TOP1 inhibitors as potent drug candidates for SARS-CoV-2-induced lethal inflammation (*8*). Therefore, we evaluated the possibility of repurposing TOP1 inhibitors to modulate neuroinflammation.

**Table 1.**
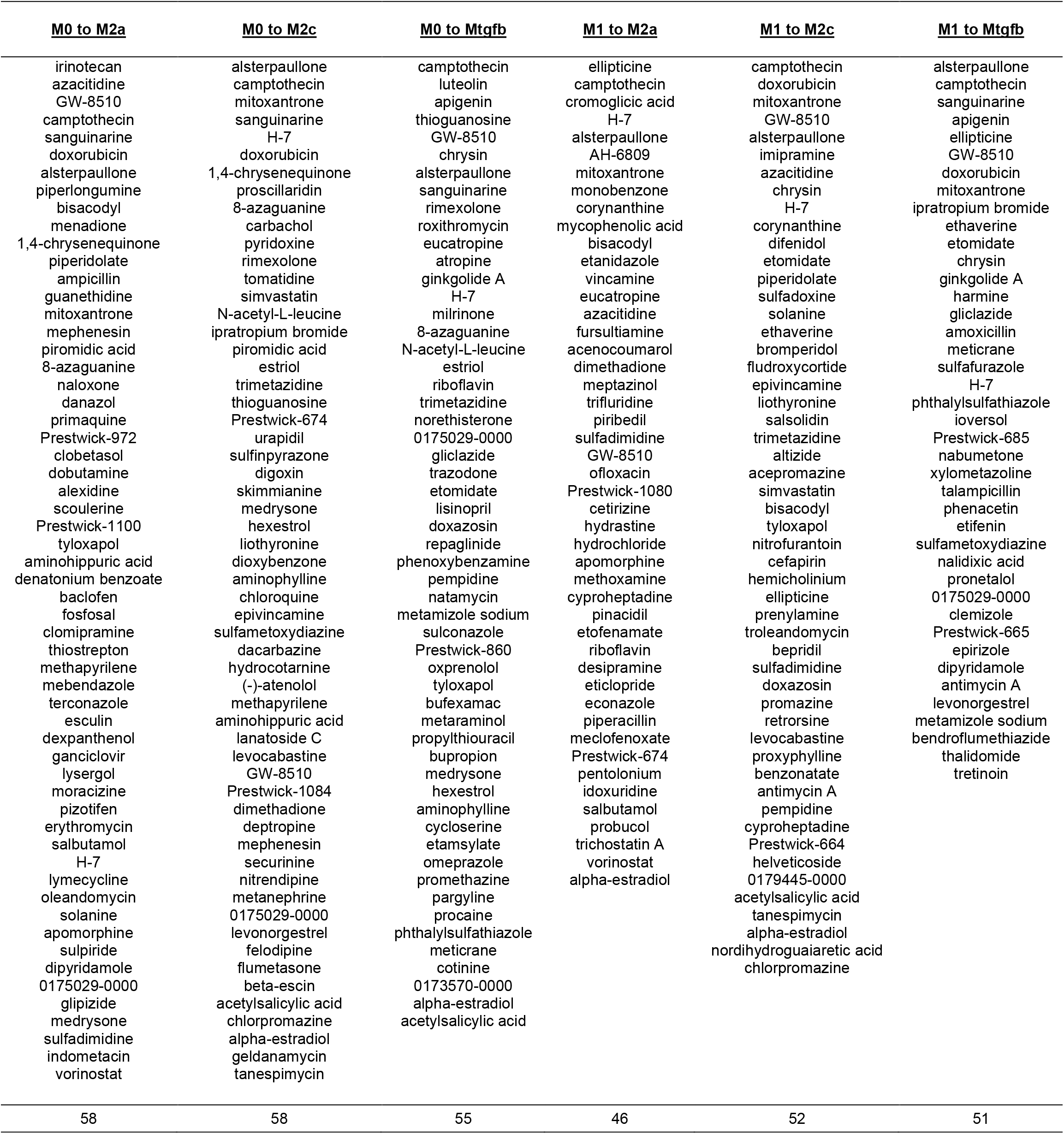
Summary of drugs and compounds in each pattern from CMAP analysis.

### TOP1 inhibition dampens inflammatory responses in stimulated microglia

We cultured primary microglia from adult mice and stimulated them with LPS/IFNγ (Fig. 2a). We observed an increased *Top1* mRNA expression after 2 h stimulation, which gradually subsided over time (Fig. 2b). The intracellular/intranuclear regulation of TOP1 activity is not fully elucidated, but some proteins have been proposed to regulate the recruitment of TOP1 to chromatin or TOP1 degradation. Among these, BTB/POZ domain-containing protein 1 (BTBD1) is known to specifically bind to TOP1, and may function as a TOP1 degradation regulator (*9*); the DNA end-processing enzyme TDP1 counteracts the biological function of TOP1 cleavage complexes (*10*); the serine/arginine-rich splicing factor 1 (SRSF1) inhibits the recruitment of TOP1 from the nucleoplasm to the nucleolus (*11*). We observed a concomitant decrease in the mRNA expression of these three negative regulators of TOP1 following microglial activation (Fig. 2c). We further confirmed an increased expression of TOP1 protein in microglia following LPS/IFNγ stimulation by Western blotting and immunostaining (Fig. 2d-f, J), indicating an active involvement of TOP1 during microglial activation.

**Fig. 2.**
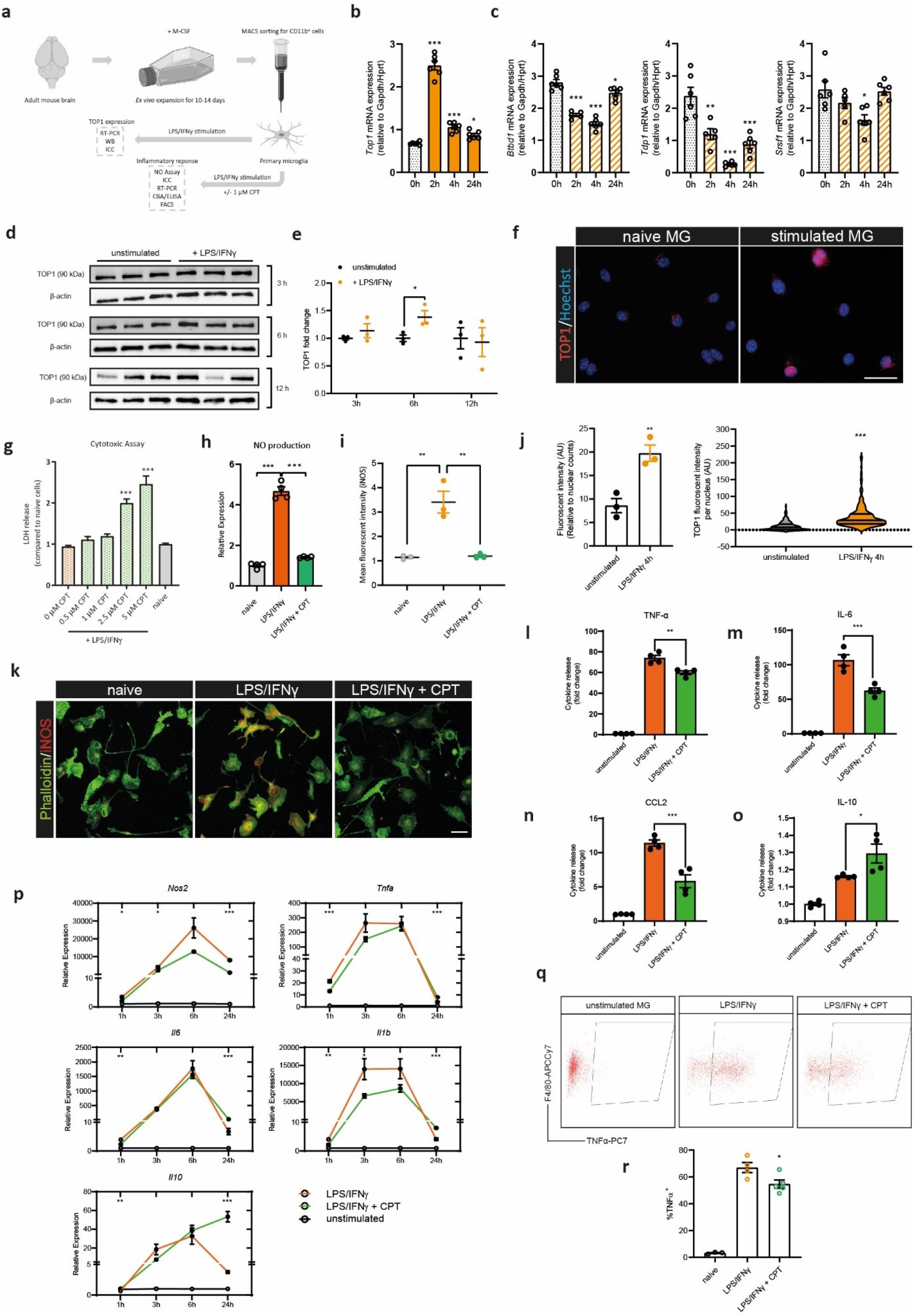
Camptothecin (CPT) impedes the inflammatory response in stimulated primary microglia. **a**, A schematic overview of the *in vitro* experiments. **b, c**, Kinetic change of mRNA expression of *Top1, Btbd1, Tdp1*, and *Srsf1* in microglia following LPS/IFNγ stimulation (*n* = 6). **d, e**, The protein expression of TOP1 was detected by Western blotting (*n* = 3); the experiment was repeated twice. **f, j**, Immunostaining of TOP1 (red) in stimulated microglia (4 h after LPS/IFNγ stimulation) or resting microglia, co-stained with Hoechst (blue) (bar = 20 μm); for the lower-left panel in (**j**), *n* = 3 per group (2 to 5 random fields per well were analyzed); for the lower right panel, 201 cells from unstimulated microglia and 256 cells from stimulated microglia were analyzed. **g**, Cytotoxicity of CPT on stimulated microglia was determined by measuring the release of LDH in the supernatants (*n* = 4). **h**, Supernatants were collected after 24 h LPS/IFNγ stimulation and measured for nitrite, as an index of nitric oxide (NO) production (*n* = 4). The graph is representative of more than three independent experiments. **i, j**, Immunostaining of iNOS (red) and Phalloidin (green) in microglia (*n* = 3 with more than 5 random fields per well) (bar = 50 μm). **l**-**o**, Cytokine release (TNF-α, IL-6, CCL2, and IL-10) to the supernatants were measured by cytometric beads assay after 24 h LPS/IFNγ stimulation (*n* = 4). The graphs show representative results of three independent experiments. **p**, The kinetics of mRNA expression of *Nos2, Tnfa, Il1b, Il6*, and *Il10* after 1, 3, 6, and 24 h of LPS/IFNγ stimulation (*n* = 3 to 6). The graphs show representative results of three independent experiments. **q, r**, The production of TNF-α within microglia was determined by flow cytometry (*n* = 3 to 5). The graphs are representative of two independent experiments. Data shown are mean ± s.e.m. \**P*<0.05, \*\**P*<0.01, \*\*\**P*<0.001. Statistical analyses were performed using two-way ANOVA with Dunnett’s Multiple Comparison test for **b** and **c**, Student’s unpaired two-tailed *t-*test for **e** and **j**, and one-way ANOVA with Dunnett’s Multiple Comparison test for the rest.

To evaluate the anti-inflammatory effect of CPT we first investigated the cytotoxicity of CPT on inflammatory microglia and a concentration below 1 μM showed no deleterious effect (Fig. 2g). We therefore used 1 μM CPT for subsequent experiments. Proinflammatory activated microglia release cytotoxic molecules such as proinflammatory cytokines and produce nitric oxide (NO) via the catalytic activity of inducible nitric oxide synthase (iNOS), which can damage CNS neural tissue (*12*). CPT incubation significantly inhibited the release of NO from stimulated microglia (Fig. 2h), which was also confirmed by a decreased iNOS immunostaining (Fig. 2i,k). Following CPT incubation the mRNA expression of inflammatory genes *Nos2, Tnfa*, and *Il1b* displayed a kinetic change; *Il6* was inhibited by CPT at 1 h, but the kinetic expression pattern was similar to that of the control group (Fig. 2p). To our surprise, the mRNA expression of the anti-inflammatory cytokine IL-10, which is important for inflammation resolution and maintenance of microglial homeostasis, was highly upregulated after 24 h in CPT-treated LPS/IFNγ-stimulated microglia.

We then measured the cytokine release in the cell supernatant from activated microglia after 24 h. The production of TNF-α, IL-6, and the monocyte chemoattractant protein CCL2 was reduced in CPT-treated microglia, whereas IL-10 production was increased (Fig. 2l-o). An increased intracellular TNF-α synthesis in stimulated microglia was also confirmed by cytokine staining using flow cytometry, and CPT treatment also inhibited the synthesis of TNF-α (Fig. 2q,r). Taken together, TOP1 is upregulated upon microglial activation, and inhibiting TOP1 dampens microglial inflammatory responses.

### The FDA-approved TOP1 inhibitor topotecan ameliorates LPS-induced neuroinflammation

Given the effect of CPT on dampening microglia inflammatory responses *in vitro*, we next assessed the effect of TOP1 inhibition *in vivo*. Due to the poor solubility of CPT, we used topotecan (TPT), which is a water-soluble analog of CPT, for *in vivo* experiments. TPT was approved by the FDA in 2007 for the treatment of small-cell lung cancer via oral administration. Similar to CPT, TPT treatment also mitigated inflammatory responses in macrophages and microglia *in vitro* (Extended Data Fig. 1).

We first employed the LPS challenge model, as it efficiently stimulates an innate immune response characterized by the activation of myeloid cells. It has been reported that one systemic LPS challenge induces robust microglial activation, depressive sickness behavior, and persistent neuroinflammation, even when peripheral immune activation is tempered (*13*). We delivered 10 μL TPT (0.1 mg/kg) directly into the CNS via intracisternal injection, followed by 5 mg/kg (i.p) LPS challenge, and analyzed the brain tissue after 4 h or 24 h (Fig. 3a). The expression of Iba1, whose immunoreactivity increases following microglial activation, was enhanced in different brain regions upon LPS treatment, whereas TPT-treated mice exhibited a reduced Iba1 immunoreactivity (Fig. 3b-d).

**Fig. 3.**
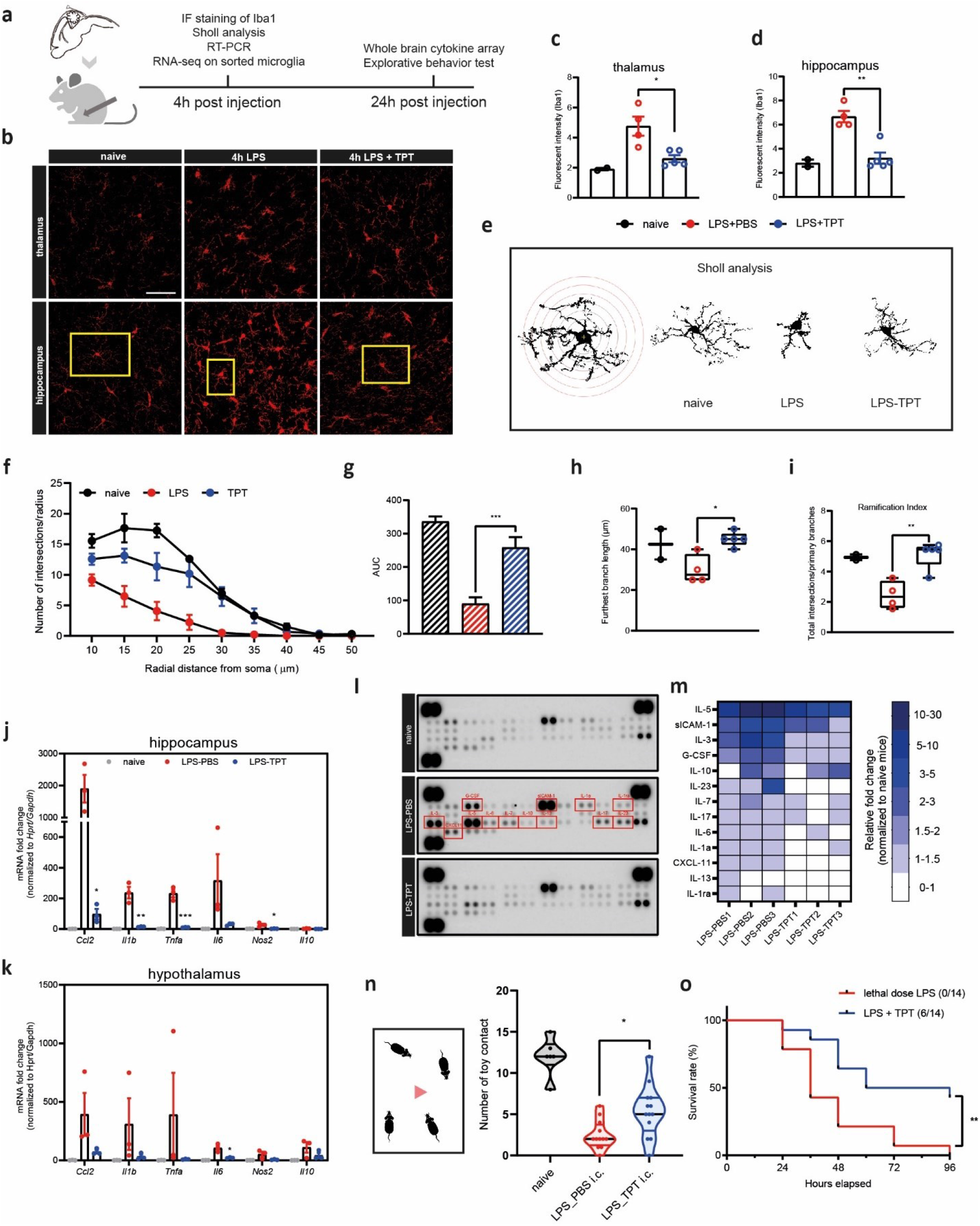
Topotecan (TPT) mitigates microglial activation and CNS inflammation in LPS-challenged mice. **a**, An overview of the experimental design. **b**, Microglia in hippocampus and thalamus were stained with Iba1 (red). **c, d**, Fluorescent intensity of Iba1 in hippocampus and thalamus; *n* = 2, 4, 5 mice for naive, LPS, and LPS + TPT group, respectively (2-6 sections per mouse were analyzed) (bar = 50 μm); representative of two independent experiments. **e**-**i**, Sholl analysis was performed to analyze the morphology of microglia. Microglia marked in yellow boxes in (**b**) were presented accordingly in (**e**); the morphological changes of microglia were objectified by the number of intersections per radius (**f** and **g**), furthest branch length (**h**), and the ramification index presented as total intersections/primary branches (**i**); *n* = 2, 4, 5 mice for naive, LPS, and LPS + TPT group, respectively (4 to 7 complete microglia from each mouse were included for analysis). **j, k**, Expression of inflammatory genes in the hippocampus and hypothalamus 4 h after the LPS challenge (*n* = 3); data are representative of two independent experiments with similar results. **l, m**, Cytokine array on brain homogenates after 24 h of LPS challenge (*n* = 2 for naive, *n* = 3 for LPS and LPS + TPT group). **n**, The contact frequency of each mouse to the new toy during the cage explorative experiment was recorded in a 5-min recording (*n* = 5, 12, 13 mice for naive, LPS_PBS, and LPS_TPT group, respectively); data are pooled from two independent experiments with similar results. **o**, A lethal dose of LPS (18 mg/kg; i.p) was given to mice; mice in LPS + TPT group received 1 mg/kg TPT 1 h before the lethal dose LPS challenge, whereas the control group received PBS injection (*n* = 14). Data shown are mean ± s.e.m. \**P* < 0.05, \*\**P* < 0.01, \*\*\**P* < 0.001. Statistical analyses comparing the LPS_PBS group with the LPS_TPT group were performed using Student’s unpaired two-tailed *t-*test; the survival curve was analyzed with Log-rank.

Microglia rapidly adapt their morphology following activation. Resting microglia are ramified cells with many intersections connecting a complex branching network, whereas activated microglia retract their long processes and become amoeboid-like and less ramified (*14*). We analyzed the morphological status of microglia using Sholl analysis (*15*), and as expected, LPS challenged mice had a typical amoeboid-like microglia morphology, while those receiving TPT treatment had an intermediate microglial morphology with increased intersections, more ramifications, and longer processes (Fig. 3e-i), indicating a less activated state. This was also reflected by a reduction in the mRNA expression of inflammatory genes in the hippocampus and hypothalamus (Fig. 3j,k). TPT treatment also resulted in a general reduction of proinflammatory cytokine levels compared to the LPS control group as evaluated by cytokine arrays of the brain homogenates (Fig. 3l,m). In addition, we also noted a better performance in an explorative behavioral test following TPT treatment (Fig. 3n), reflecting less depressive-like sickness. Meanwhile, we also injected 10 ng of LPS directly into the CNS via intracisternal injection to limit inflammation within the CNS and administered 1 mg/kg TPT (i.p). In this experimental setting, we also observed an improved behavior following TPT treatment compared to the LPS control group (Extended Data Fig. 2). A previous study reported that CPT rescued mice from a lethal LPS challenge (*16*). Similarly, we determined that a single low dose injection of TPT (1 mg/kg) could also prevent around 40% of mice from death induced by lethal inflammation (Fig. 3o). Together, these results confirmed the protective effect of TPT on neuroinflammation induced by the LPS challenge.

### TPT drives an altered transcriptional phenotype in microglia of LPS-challenged mice

The beneficial effect of TPT in the LPS model prompted us to further characterize the cellular and molecular responses in microglia following TPT treatment. To understand the molecular changes, we sorted microglia 4 h after the LPS challenge and performed RNA sequencing. The multidimensional scaling (MDS) analysis clearly separated the naive control from the LPS (LPS + PBS) and TPT (LPS + TPT) group (Fig. 4a). We observed that microglial homeostatic genes (*P2ry12, Tmem119, Siglech, Olfml3*) were significantly downregulated, indicating an immediate microglial activation after the LPS challenge (Extended Data Fig. 3a). As expected, we confirmed molecular signatures (IL-17, MAPK, NF-κB, TNF, and Toll-like receptor signaling pathways) indicative of activated inflammatory responses in microglia of LPS-challenged mice (Extended Data Fig. 3b-d).

**Fig. 4.**
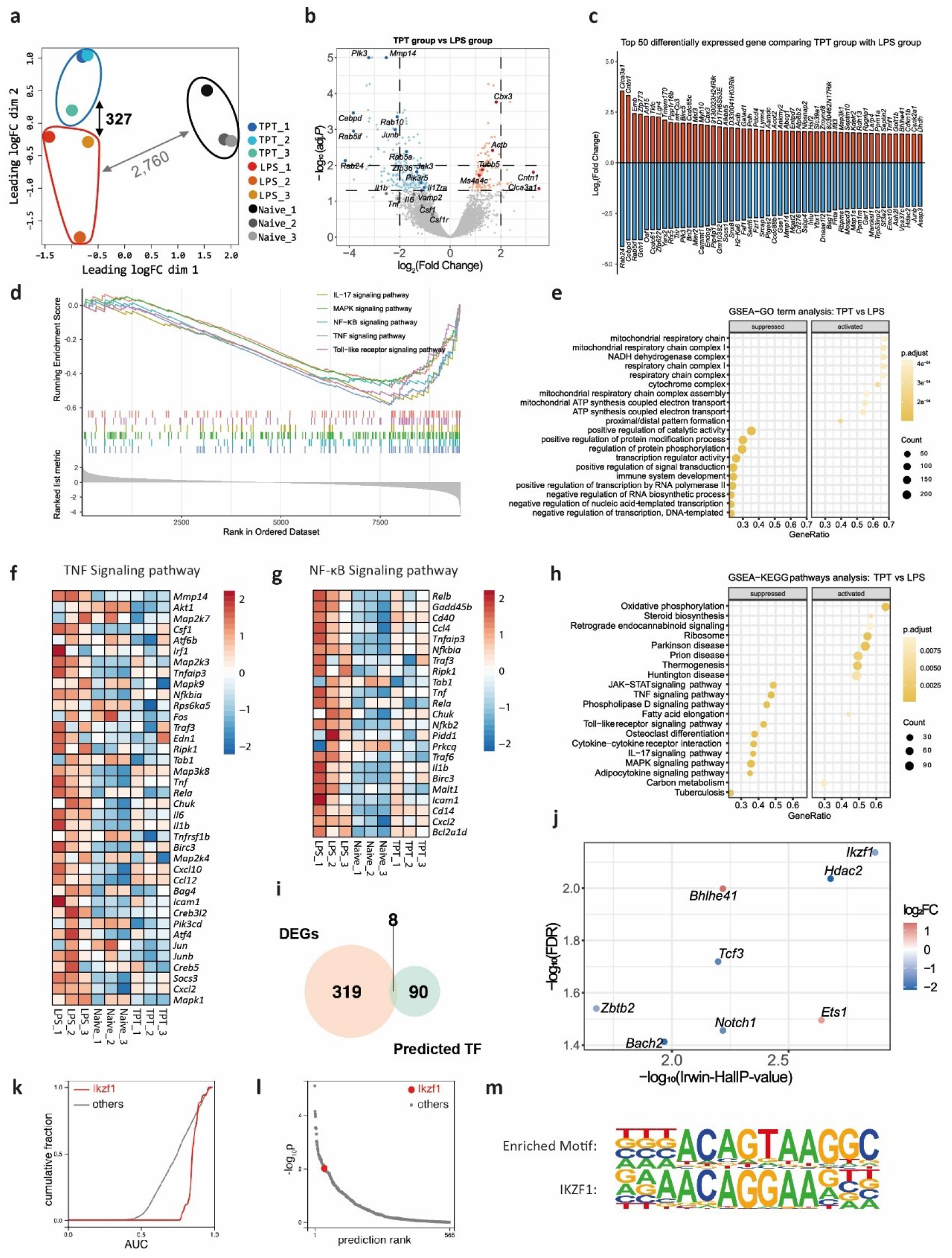
Topotecan reshapes the transcriptional profile of microglia after systemic LPS challenge. **a**, Multidimensional scaling (MDS) analysis of brain microglia sorted from the following groups (*n* = 3): naive group (mice received i.p injection of PBS and intracisternal injection of PBS), LPS group (mice received i.p injection of 5 mg/kg LPS following intracisternal injection of PBS), and TPT group (mice received i.p injection of 5 mg/kg LPS following intracisternal injection of 0.1 mg/kg TPT); numbers labelled between different clusters indicate the number of differentially expressed genes (DEGs) identified by EdgeR (FDR=<0.05). **b**, Volcano plot depicting the DEGs between the TPT and LPS groups; dots represent non-DEGs (grey, FDR>0.05), DEGs with fold change log_2_FC> 0 (red), and DEGs with log_2_FC<0 (blue); genes of interest were labelled and plotted in darker colors. **c**, Bar plot of the top 50 up-and downregulated DEGs (ordered by FC) between the TPT and LPS groups. **d**, GSEA plot of the enriched KEGG pathways related to inflammation between the TPT and LPS groups. **e, h**, Dot plots showing the top10 suppressed and activated GO terms (**e**) or KEGG pathways (**h**) between the TPT and LPS groups. **f, g**, Heatmaps plotting the changes in expression levels of genes contributing to the enrichment analysis of TNF signaling pathway (**f**) and NF-κB signaling pathway (**g**) across samples when comparing the TPT group with the LPS group. The expression levels were calculated by TPM and log2 transformed. **i**, Venn diagrams intersected the number of DEGs and transcription factor (TF) predicted to regulate the gene expression between the TPT and LPS groups. **j**, The Scatter plot shows the Irwin-Hall and FDR value of the 8 genes in (**i**); FDR was obtained from differential expression (DE) analysis; Irwin-Hall *P*-value was obtained from BART prediction, indicating the integrative ranking significance; the color gradient indicates FC in DE analysis. **k**, BART output of association score cumulative distribution of Ikzf1. **l**, BART output of transcriptional regulator prediction rank of Ikzf1. **m**, *De novo* motif enriched in promoters of DEGs between the TPT and LPS group matching to known motifs of Ikzf1 using HOMER.

We next examined differences in gene expression between the TPT and LPS groups. Although LPS caused global gene expression changes in microglia, the TPT group was still distinct from the LPS group according to the MDS analysis, with 327 differentially expressed genes identified (Fig. 4a). We noted that the inflammatory pathways activated following the LPS challenge were significantly inhibited in the TPT group, including TNF and NK-κB signaling pathways, which further corroborated our findings of a potent anti-inflammatory and protective effect of TPT in activated microglia and LPS challenged mice (Fig. 4d,f,g; Extended Data Fig. 3e-h).

Among the top 50 downregulated genes, many are key regulators controlling inflammatory responses in microglia, such as *Cebpd, Mmp14, Hdac2*, and *Junb* (Fig. 4b,c). The expression of *Cebpb, Mmp14, and Hdac2* was also significantly increased in the LPS group when compared to the naive group (Extended Data Fig. 3a). Of note, JunB is one of the three JUN family members (together with c-Jun and JunD) and a subunit of the transcription factor complex containing activator protein 1 (AP-1). The JunB/AP-1 complex acts downstream of several inflammatory signaling pathways, including MAPK and TLR, that regulate gene expression in response to stimuli such as stress and cytokines (*17, 18*). As a central transcription factor, CEBPD is essential for regulating proinflammatory gene expression and microglial activation (*19, 20*). We also noted that several small GTPase genes (*Rab5a, Rab5if, Rab10, Rab24*) were significantly downregulated. Many of these genes are involved in endosomal pathways and intracellular vesicle trafficking, which are processes important for cytokine transportation and release.

In addition to suppression of many inflammatory pathways following TPT treatment, transcripts of oxidative phosphorylation, steroid biosynthesis, and mitochondrial respiratory activities were enriched, indicating a potential metabolic reprogramming in microglia (Fig. 4e,h). We also predicted the potential transcription factors (TFs) mediating TPT effects by using Binding Analysis for Regulation of Transcription (BART) and Hypergeometric Optimization of Motif EnRichment (HOMER) analysis. From the ChIP-seq data-based BART analysis we obtained 90 predicted TFs, and among those, eight were differentially expressed when comparing TPT and LPS groups (*Zbtb2, Tcf3, Notch1, Ikzf1, Hdac2, Ets1, Bhlhe41, Bach2*) (Fig. 4i,j; Extended Data Fig. 3i). By combining the confidence of transcription factor prediction (Irwin-Hall p-value) with differential expression analysis, *Ikzf1* and *Hdac2* ranked the highest (Fig. 4k,l; Extended Data Fig. 3i). Furthermore, the known motif of *Ikzf1* showed high similarity to one of the top-ranked motifs that occurred in the promoter regions of differentially expressed genes (Fig. 4m; Extended Data Fig. 3j). This indicates that TPT may protect microglia against inflammatory activation by downregulating *Ikzf1* which commonly regulates pro-inflammatory gene transcription, together with *Hdac2* (*21*–*23*). Taken together, our results reveal a less inflammatory transcriptional phenotype of microglia following TPT treatment with key proinflammatory regulators being remodeled.

### TPT prevents disease progression in experimental autoimmune encephalomyelitis

We next employed the experimental autoimmune encephalomyelitis (EAE) model for investigation. Unlike the LPS model in which myeloid cells are the main effector cells, EAE features activated Th1/Th17 cell infiltration into the CNS together with proinflammatory myeloid cells, causing devastating demyelination reflected by clinical motor symptoms (*24*). Thus, EAE mimics the immunopathogenesis of MS to a certain extent. For testing the effect of TPT in EAE we employed a treatment protocol comprising 3 injections of TPT (1 mg/kg; i.p) (Fig. 5a). This is a much lower dose (5 to 10-fold reduction) compared to previous studies using TPT as a typical chemotherapeutic agent in rodent models and is equivalent to the well-tolerated dose reported in a mouse model of Huntington’s disease (*25*–*27*). We characterized that this treatment protocol did not induce loss of body weight, nor a significant decrease in the number of peripheral immune cells, excluding the unwanted immunosuppression and cytotoxic effects (Extended Data Fig. 4).

**Fig. 5.**
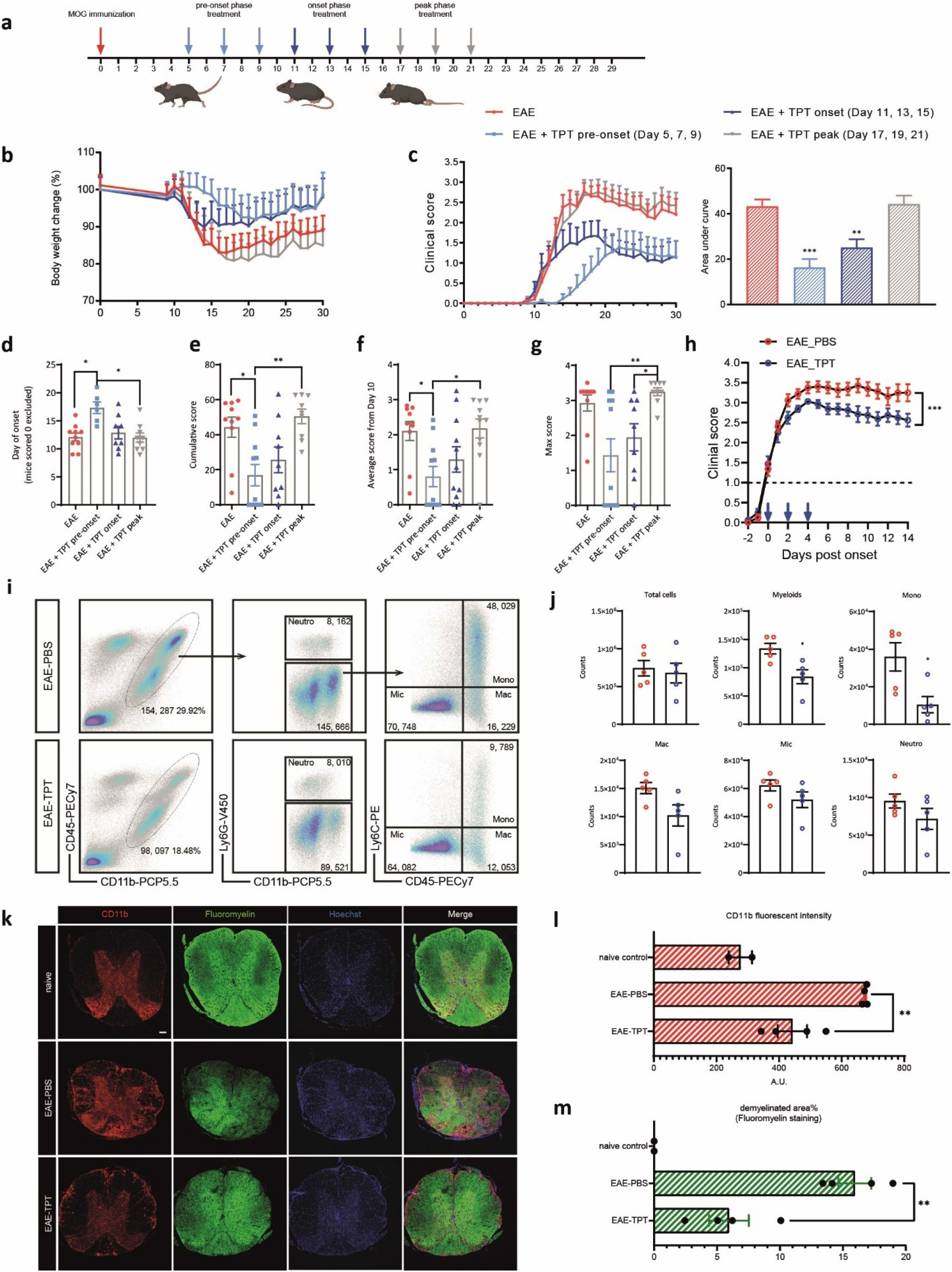
Topotecan prevents disease progression in the EAE model. **a**, Mice received TPT treatment (1 mg/kg, i.p) at indicated days post-immunization: pre-onset group (day 5, 7, 9), onset group (day 11, 13, 15), and peak group (day 17, 19, 21); mice in the EAE control group received 3 PBS injections randomly from day 5 (i.p). **b**, Bodyweight change following EAE immunization. **c**-**g**, EAE clinical scores among the groups were analyzed using different parameters: area under the curve (**c**), day of onset (**d**), cumulative score (**e**), the average score from day 10 (**f**), and max score (**g**); *n* = 9 -11 per group; data are from one EAE batch representing three batches with a similar effect. **h**, EAE mice were treated individually from the day when they were just scored or scored higher than 1; *n* = 8 mice per group. Data were representative of two independent batches. **i**, EAE mice received TPT treatment from the onset or PBS were euthanized at the peak of EAE (day 18), and the spinal cords were dissected for flow cytometry with an antibody panel for myeloid cells (*n* = 5), and the counts of different myeloid cells, including monocytes (Mono), macrophages (Mac), microglia (Mic) and neutrophils (Neutro), were graphed in (**j**); this experiment was performed twice. **k**, Spinal cord sections dissected from Day 30 from the EAE batch in (**h**) were stained with CD11b (red), Fluoromyelin (green), and Hoechst (blue) (bar = 100 μm); the CD11b immunofluorescent intensity was graphed in (**l**), and the percentage of demyelinated area in the white matter of the spinal cord was graphed in (**m**). Data are presented as the mean ± s.e.m. \**P* < 0.05; \*\**P* < 0.01; \*\*\**P* < 0.001. The following statistical analyses were used: the EAE scores in (**d**-**g**) were analyzed using the Kruskal-Wallis nonparametric test; the AUC in (**c**) was analyzed using the one-way ANOVA with Dunnett’s Multiple Comparison test, and the Student’s unpaired two-tailed *t-*test was used for comparing the EAE-PBS group with the EAE-TPT group in **h, j, l**, and **m**.

We included three TPT treatment groups with: (i) the *pre-onset* group receiving TPT on days 5, 7, and 9 post-immunization; (ii) the *onset* group receiving TPT on days 11, 13, and 15; and (iii) the *peak* group on days 17, 19 and 21. EAE mice gradually lost weight upon disease onset, but mice in the TPT pre-onset and onset treatment groups experienced less weight loss (Fig. 5b). TPT pre-onset treatment significantly delayed EAE disease onset and prevented some mice from developing the disease (Fig. 5c-g). Both TPT pre-onset and onset treatment improved the EAE clinical score (Fig. 5c), while TPT treatment starting from disease peak failed to do so. We also dissected the spinal cords at days 18 and 30 and confirmed a reduced infiltration of peripheral monocytes into the spinal cord at EAE peak, and a smaller demyelinated area at EAE recovery phase following TPT onset treatment (Fig. 5i-m). Furthermore, we analyzed the functionality of the immune cells in the spinal cords at EAE peak from the TPT onset treatment group and PBS control group. There was no difference in the frequency of Ki67+ CD4+ proliferating T cells, as well as Foxp3+ CD4+ regulatory T cells (Extended Data Fig.5c,d). In terms of cytokine production, we did not observe a difference in GM-CSF and IFN-γ production of CD4+ T cells between the two groups (Extended Data Fig.5e,f), but we noted a significantly increased production of IL-10 in CD11b+ myeloid cells in TPT-treated mice, while the difference in TNF-α production was not significant (Extended Data Fig.5a,b). Since the day-of-onset for each EAE mouse varied even within the same EAE batch, we then treated the EAE mice individually from the day the mouse was scored 1 for the first time, thereby ensuring that all mice were sick and at an early onset phase when treatment commenced (Fig. 5h). We again observed a significant therapeutic effect of TPT in this experimental setting. These results, together with the effect of TPT in the LPS challenge model, led us to conclude that TPT could be repurposed for the treatment of neuroinflammatory conditions.

### TOP1 expression is elevated in neuroinflammatory conditions

Being the target of TPT we next addressed whether TOP1 levels were altered during neuroinflammatory conditions (Fig. 6a).

**Fig. 6.**
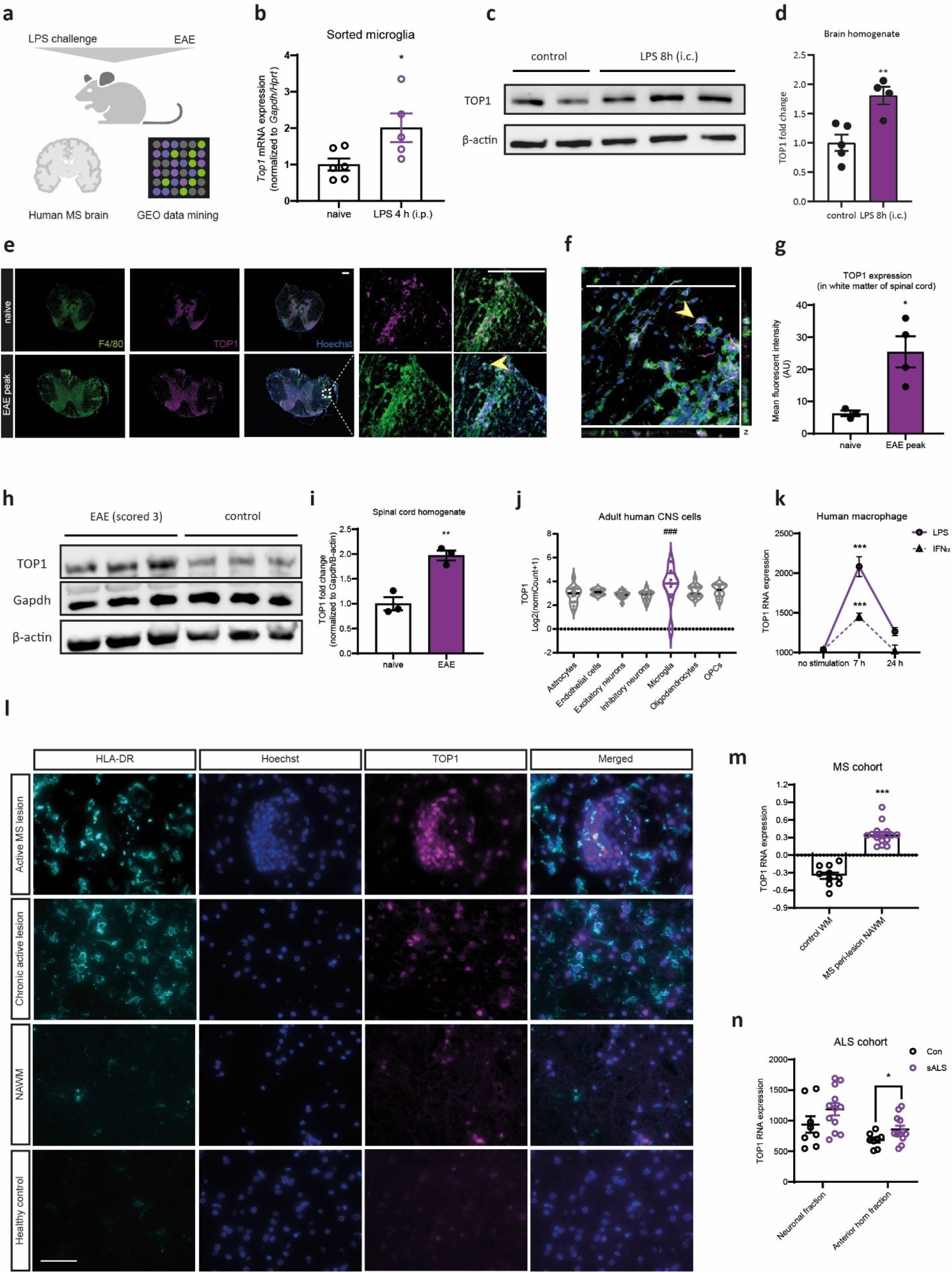
TOP1 expression is increased in neuroinflammatory conditions. **a**, Expression of TOP1 was examined in the LPS and EAE mouse models, and MS brain sections; public datasets were also exploited to ascertain TOP1 expression. **b**, Microglia were sorted from LPS challenged mice (i.p) after 4 h or naive mice, and the mRNA expression of TOP1 was graphed (*n* = 5 or 6); the experiments were performed twice. **c, d**, Western blotting of TOP1 expression from brain homogenates of mice receiving intracisternal LPS injection or PBS after 8 h (*n* = 4); graphs are representative of two experiments. **e**-**g**, Immunofluorescent staining of spinal cord sections from mice at EAE peak and control mice with anti-F4/80 (green), TOP1 (magenta), and nuclei with Hoechst (blue); a magnified image of the selected region in (**e**) is shown in (**f**) and the mean fluorescent intensity of TOP1 in the white matter was graphed in (**g**); *n* = 3 to 4 mice per group with 3-5 sections from each mouse; bar = 200 μm; the yellow arrow indicates a representative F4/80+ cell with increased TOP1 expression in the nucleus. **h**, TOP1 expression in the spinal cord homogenates of EAE mice dissected at disease peak (scored 3) or control mice was examined by Western blotting, and quantified in (**i**) with normalization to GAPDH and β-actin (*n* = 3). **j**, TOP1 expression in healthy adult human CNS cells was extracted from the Brain Myeloid Landscape plotted with the dataset GSE97930. **k**, Data mining on TOP1 expression from the dataset GSE85333. **l**, Brain sections from healthy control and MS brain tissues, including the active MS lesion, chronic active lesion, and the normal-appearing white matter (NAWM), were stained with anti-HLA-DR, anti-TOP1, and co-stained with Hoechst (bar = 50 μm). **m, n**, Data mining on TOP1 expression from GSE108000 (an MS cohort containing NAWM from 15 MS cases, and white matter from 10 healthy controls) and GSE18920 (an ALS cohort containing gene expression data from neuronal fraction and anterior horn fraction of 8 healthy donors and 12 sporadic ALS patients). Data are presented as the mean ± s.e.m. \**P* < 0.05; \*\**P* < 0.01; \*\*\**P* < 0.001. The student’s unpaired two-tailed *t-*test was performed for statistical analyses except (**k**), which was performed using two-way ANOVA with Dunnett’s Multiple Comparison Test. ### *P*_adj_ ≤ 1e-04 when comparing microglia with other cell types, according to the analysis from the Brain Myeloid Landscape 2.

In mouse microglia sorted 4 h post-LPS challenge (i.p), we observed an upregulation of *Top1* mRNA expression (Fig. 6b). When 10 ng LPS was delivered directly into the CNS via intracisternal injection, we also noted an increase of TOP1 protein expression in the brains 8 h later (Fig. 6c,d). In parallel, we also observed an increase of TOP1 expression in the spinal cords obtained from mice at the EAE peak stage (score 3), as evidenced by both immunofluorescent staining and Western blotting (Fig. 6e-i). TOP1 is a nuclear protein, and accordingly, the enhanced TOP1 immunoreactivity was mainly evident in the nuclear compartment of F4/80-expressing myeloid cells, which represent the infiltrating monocytes/macrophages in the context of EAE (Fig. 6e,f).

By exploring public genomic datasets we found that under physiological conditions, microglia/myeloid cells have higher baseline expression of *TOP1* compared to other cell types in adult CNS (Fig. 6j; Extended Data Fig. 6). In another dataset, stimulation with LPS or IFNα for 7 h was sufficient to increase TOP1 expression in human monocyte-derived macrophages, with a return to baseline levels already 24 h post-challenge (Fig. 6k). In an MS cohort, the peri-lesion normal-appearing white matter (NAWM) had higher *TOP1* expression compared to the healthy control white matter (Fig. 6m). We further confirmed this by evaluating TOP1 expression in brain tissues from MS patients, and in a like manner, a pronounced TOP1 expression in the lesion areas as well as the NAWM, compared to the white matter from healthy control donors, was evident (Fig. 6l). In a sporadic ALS cohort, patients also showed increased *TOP1* expression in the anterior horn fraction (enriched for glial cells) compared to healthy controls, but this was not a prominent feature in the neuronal fraction (Fig. 6n). Taken together, these results indicate that increased TOP1 expression is characteristic of several neuroinflammatory conditions and that TOP1 is a potential target for controlling neuroinflammation.

### Nanoengineered DNA origami for specific delivery of TPT to myeloid cells

Even though a low dose application of TPT had little toxicity, we aimed for more specific delivery of TPT to myeloid cells. From a translational perspective, this would even enhance the effect of TPT on myeloid cells, and meanwhile, minimize the potential off-target effects on other cell types when applying TOP1 inhibition therapy in patients.

To specifically deliver TPT and to restrict its effect to myeloid cells, we took advantage of a DNA origami system to load TPT and coated it with β-glucan, which is a ligand from baker’s yeast recognized mainly by the Dectin-1 receptor expressed on myeloid cells (*28*). The superiority of using DNA origami as a drug delivery vehicle is its precision and reproducibility (*29, 30*). Unlike other self-assembled nanoparticles that often have a relatively broad size distribution, DNA origami has the same controlled shape, size, and charge for each particle. The DNA origami nanostructure used in this work was a wireframe-stylized hexagonal rod (HR) with a length of ∼120 nm and a diameter of ∼20 nm (Fig. 7a). To coat the HR with β-glucan we designed 60 evenly distributed protruding single-stranded DNAs around the HR for the hybridization of single-stranded polyA50 (at the 3 prime) containing DNA oligos. One polyA50 oligo and two glucan polymers can then form a stable triple-helical conformation at the site where polyA50 is located on HR, forming glucan-coated HR (HR-Glu) (*31, 32*). Previous studies have revealed that TPT binds to single-stranded nicks in double-stranded DNA (dsDNA) through which it stops topoisomerase to correct the overwinding or underwinding of DNA during DNA replication and transcription (*33, 34*). TPT could potentially also intercalate between the bases in dsDNA (*35*). These interactions between TPT and dsDNA thus enabled the loading of TPT into the DNA origami nanostructure without chemical conjugation (Fig. 7a).

**Fig. 7.**
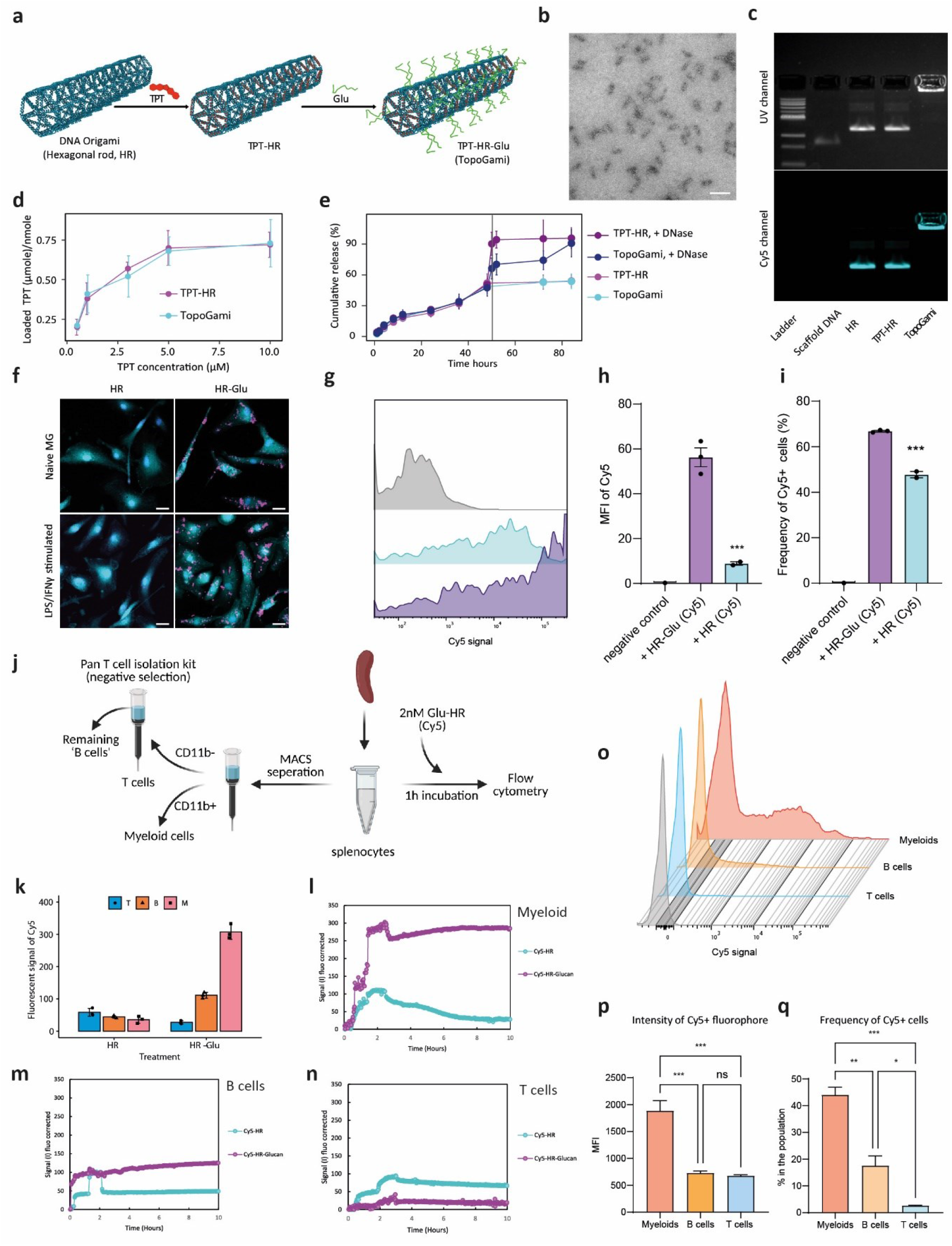
A bioengineered DNA-origami nanosystem for selective topotecan (TPT) delivery to myeloid cells. **a**, TPT was loaded into the hexagonal rod (HR)-structured DNA origami with surface modification by β-glucan. **b**, Ultrastructural confirmation of the HR structure under transmission electron microscopy (bar = 100 nm). **c**, Cy5-labeled DNA nanostructures (HR, TPT-HR, and TopoGami) electrophoresis on 2% agarose gel; the gel is pre-stained with ethidium bromide and imaged under the UV channel and Cy5 channel. **d**, TPT loading efficacy of HR and glucan-coated HR. **e**, Cumulative TPT release profiles of TPT-HR and TopoGami with or without DNase adding at the time point 48 h. **f**, Uptake of Cy5-labeled HR or HR-Glu structures (in magenta) by non-stimulated or stimulated mouse primary microglia (in cyan) after 1 h incubation with 2 nM of the structures (bar = 20 μm); the experiment was performed twice. **g**-**i**, Flow cytometry analysis comparing the uptake of Cy5-labeled HR or HR-Glu structures in non-stimulated microglia incubated with 2 nM of the structures for 1 h (*n* = 3); the quantification was based on the median fluorescent intensity (MFI) or the percentage of Cy5-expressing cells in total cells; data are representative of two independent experiments. **j**, Splenocytes were processed for comparing the uptake of Cy5-labeled HR-Glu structure in different immune cells. The cells were separated using magnetic-activated cell sorting (MACS) into 3 compartments (myeloid cells, T cells, and B cells) for subsequent LigandTracer experiments in (**k**-**n**), or directly incubated with Cy5-labeled Glu-HR for flow cytometry analysis (**o**-**q**). **k**, The fluorescent signal of Cy5 in different immune cell compartments after 8 h incubation with HR or HR-Glu in LigandTracer experiments; T for T cells, B for B cells, and M for myeloid cells; data are pooled from three observations. **l, m**, The real-time interaction of Cy5-labeled HR or HR-Glu structures to myeloid cells, B cells, and T cells. 2 nM of structures were added to the cells at 0 h, and 6 nM of structures were added to the cells at 1 h; the unbonded structures were removed after 2-3 h and the cells were kept in a fresh medium afterward. **o**-**q**, Flow cytometry analysis of Cy5 fluorescent signal in CD11b^+^CD45^+^ myeloid cells, CD11b^-^CD45^+^CD19^-^CD3^+^ T cells, and CD11b^-^CD45^+^CD19^+^CD3^-^ B cells; the quantification was based on the median fluorescent intensity (MFI) or the percentage of Cy5-expressing cells in each population (*n* = 3); the experiment was repeated twice with similar results. Data are presented as the mean ± s.e.m. \**P* < 0.05; \*\**P* < 0.01; \*\*\**P* < 0.001. The statistical analysis was performed using One-way ANOVA with Dunnett’s Multiple Comparisons Test.

We prepared the TPT-loaded and glucan-coated HR (HR-TPT-Glu) and denoted this system ‘TopoGami’. We confirmed the expected hexagonal rod structure of TopoGami by transmission electron microscopy (Fig. 7b). Agarose gel electrophoresis of Cy5-labeled HR, TPT-HR, and TopoGami revealed that all samples had sharp-structured bands, indicating the high quality and consistency of structure production irrespective of TPT loading. More importantly, TopoGami migrated poorly during electrophoresis, reflecting the fact that the structure was successfully coated by glucan as it neutralized the negatively charged DNA (Fig. 7b,c). We determined that the saturated TPT loading capacity in 1 nM of HR was ∼600 nM, and glucan coating had no effect on the loading efficiency (Fig. 7d). We also characterized the stability of the drug-loading DNA origami structure by measuring the cumulative TPT release over time. The release profile of TPT revealed no sudden increase within 84 h, indicating the stable integration between TPT and the DNA nanostructure. When we added DNase, we observed that TPT release from TopoGami accelerated gradually, whereas its release from the TPT-HR DNA origami without glucan coating surged immediately (Fig. 7e). This differential sensitivity to DNase thus suggested that β-glucan coating, in addition to acting as a myeloid-binding ligand, can also protect the DNA nanostructure from protein accessibility and prevent its degradation.

To confirm glucan-mediated internalization of the DNA structure, we incubated mouse primary microglia with 2 nM HR or HR-Glu in both resting and activated states for 1 h. HR-Glu was efficiently taken up by microglia compared to HR, irrespective of microglial activation state (Fig. 7f). This was further demonstrated by flow cytometry as an increased Cy5 fluorophore intensity was observed in microglia treated with HR-Glu compared to those exposed to HR (Fig. 7g-i).

We next assessed the interaction of glucan-coated HR to different immune cells using splenocytes as a source. We separated the splenocytes into three major immune compartments (myeloid cells, T cells, and B cells) and performed real-time structure-to-cell interaction assay with each compartment (Fig. 7j). All the three cell compartments had limited interactions with the non-coated HR (Cy5-labelled), but the interaction between the Cy5-labelled HR-Glu and myeloid cells was much higher than with B and T cells (Fig. 7k). During the real-time measurement, HR-Glu had a faster association with myeloid cells (Fig. 7l) and a slight association with B cells (Fig. 7m) compared to HR; however, little association of HR-Glu was evident with T cells (Fig. 7n). To further validate these findings, we incubated splenocytes with Cy5-labelled HR-Glu and performed flow cytometry (Fig. 7j). We gated for the 3 cell types (Extended Data Fig. 7) and again observed a significantly increased uptake of the HR-Glu DNA origami in the myeloid compartment (Fig. 7o-q). These results indicate that the glucan-coated HR DNA origami is a reliable system to deliver TPT to myeloid cells.

### TopoGami recapitulates the anti-inflammatory effects of TPT

We next verified the anti-inflammatory properties of TopoGami. We harvested the cells after 4 h stimulation and performed RT-PCR analysis of inflammatory genes. Compared to the control group, TopoGami strongly inhibited the mRNA expression of *Nos2, Il1b, Il6*, and *Tnfa* (Fig. 8a-d). We also analyzed cytokine release in the supernatants from treated cells after 24 h, and TopoGami significantly inhibited the release of IL-6 and reduced TNF-α production to a large extent (Fig. 8e,f). To our surprise, we noted that HR-Glu itself also exhibited a certain degree of anti-inflammatory properties. However, with the addition of TPT, TopoGami had a greater inhibitory action.

**Fig. 8.**
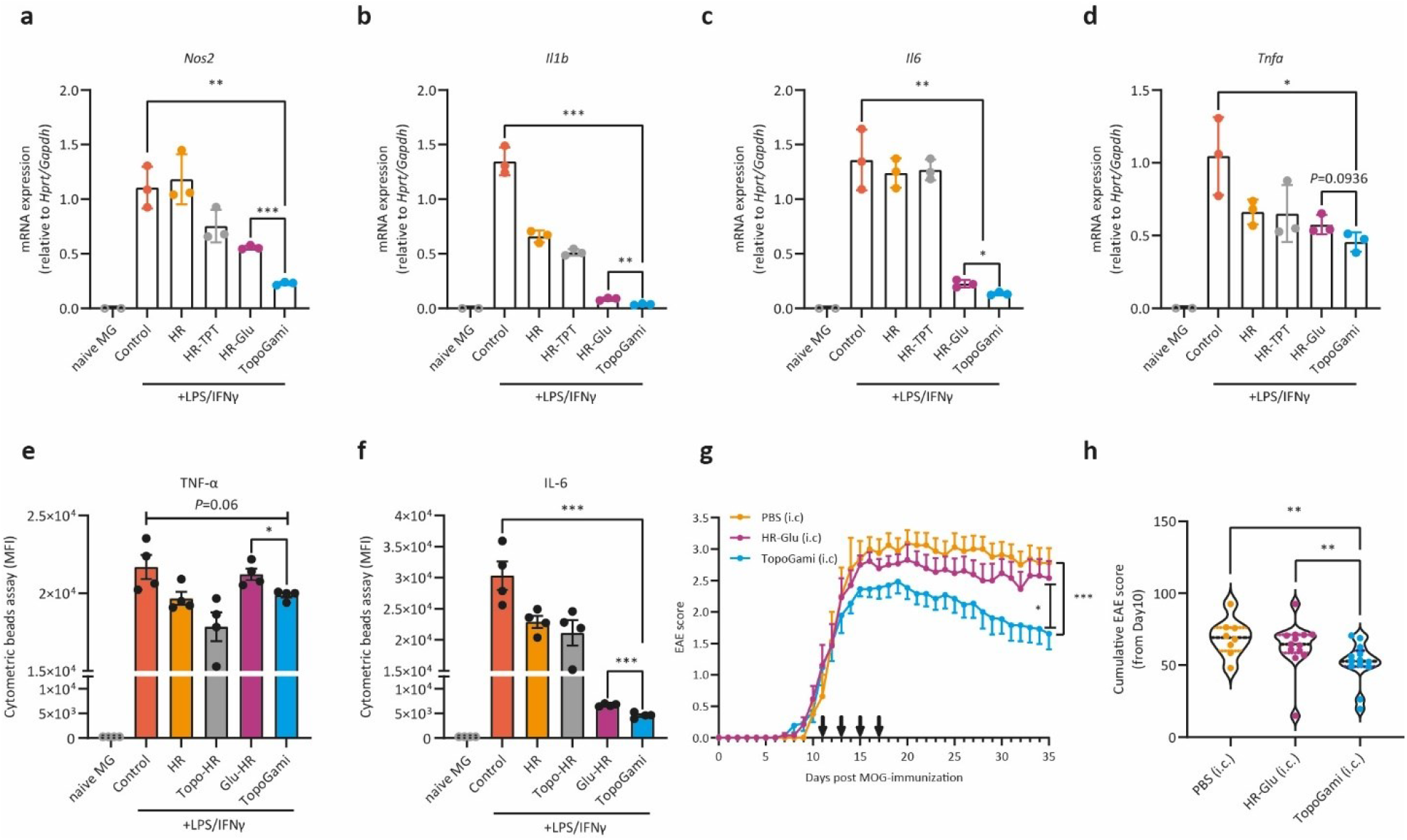
TopoGami inhibits the inflammatory responses in stimulated microglia and ameliorates EAE. **a**-**d**, The mRNA expression of *Nos2, Il1b, Il6*, and *Tnfa* in microglia treated with HR, TPT-HR, HR-Glu, or TopoGami after 4 h LPS/IFNγ stimulation (*n* = 3); data are representative of two independent experiments. **e, f**, The cytokine release of IL-6 and TNF-α from microglia treated with HR, TPT-HR, HR-Glu, or TopoGami after 24 h LPS/IFNγ stimulation were measured by cytometric beads assay (*n* = 4); the experiment has been repeated twice. **g, h**, Mice with EAE received intracisternal injections of TopoGami, HR-Glu, or PBS, at days 11, 13, 15 17 post-immunization (*n* = 8 for the PBS group; *n* = 13 for the TopoGami and HR-Glu groups); data are pooled from two independent EAE batches. Data are presented as the mean ± s.e.m. \**P* < 0.05; \*\**P* < 0.01; \*\*\**P* < 0.001. The comparison between the TopoGami group and control group, or the TopoGami group and Glu-HR in (**a**-**f**) was performed using the Student’s unpaired two-tailed *t-*test; the analysis of the AUC in (**g**) was analyzed using one-way ANOVA with Dunnett’s Multiple Comparison test; Kruskal-Wallis nonparametric test was performed for EAE cumulative score in (**h**).

To assess the ability of TopoGami to ameliorate neuroinflammation, we employed the EAE model. We first administered TopoGami with 100 µM loaded TPT (100 μL per mouse) via i.p injection at days 5, 7, and 9 after immunization as a prophylactic treatment. Similar to results with TPT itself, i.p injection of TopoGami also delayed EAE progression (Extended Data Fig. 8). We then assessed if targeting microglia and infiltrating inflammatory myeloid cells in the CNS alone could confer beneficial effects. Therefore, we delivered 10 µL TopoGami directly into the CNS via intracisternal injection from the EAE onset phase (injected at days 11, 13, 15, and 17 post-immunization). TopoGami treatment significantly improved the clinical symptoms compared to both the PBS group and the HR-Glu vehicle control group (Fig. 8g,h). These results demonstrate that targeting myeloid cells using TopoGami mitigates microglial activation and neuroinflammation.

## Discussion

The importance of myeloid cells in neuroinflammatory conditions has been acknowledged greatly during recent years. Other than depleting myeloid cells using CSF1R inhibitors, preclinical therapeutics regulating myeloid cell function are scarce. Based on this status quo, the purpose of our study was to discover novel therapeutics that regulate dysfunctional myeloid cells (particularly microglia) and to develop practical drug delivery tools to potentiate the specific targeting of myeloid cells. Several landmark studies have highlighted TOP1 inhibition as a strategy for virus infection-induced inflammation, with a recent study reporting that TPT suppresses lethal inflammation induced by SARS-CoV-2 in hamsters (*16, 36*). Serendipitously, we independently identified TOP1 inhibitors as potent anti-inflammatory agents for inflammatory microglia and demonstrated their efficacy in treating neuroinflammatory diseases. Importantly, we designed a novel nanoengineered TOP1 inhibitor (TopoGami) that could efficiently and specifically target myeloid cells and reduce their proinflammatory activation state.

We screened a list of compounds with underlying microglia-modulating function using *Connectivity Map*. Apart from camptothecin, the high rank of mitoxantrone on the list is also noteworthy. Mitoxantrone was originally developed as an antineoplastic drug but was repurposed and approved for use in relapsing-remitting MS and aggressive progressive MS (*37, 38*). Although the mechanism of mitoxantrone in treating MS was not fully elucidated when it was approved, it has been determined to inhibit microglial inflammatory responses by antagonizing TLR4 and suppressing downstream NF-κB activation (*39*). Not only does this validate our screening strategy, but our screened drug lists also provide insights for the development of future therapeutics for neuroinflammation.

Although the precise mechanism of why peripheral LPS challenge activates microglia in the CNS is not fully understood, neuroinflammation induced by systemic LPS challenge represents a useful proof-of-concept model to study microglial function. LPS might access the brain parenchymal through lipoprotein-mediated transport and bind to TLR4 expressed abundantly on microglia, and at a dose higher than 3 mg/kg LPS also disrupts the blood-brain barrier (*40, 41*). A key study proposed that systemic LPS-induced microglial activation is mediated via TNF-α production in the liver and its subsequent translocation to TNF receptors on microglia, inducing downstream NF-κB signaling cascades (*13*). Of note, this study showed that a single systemic LPS challenge (5 mg/kg) could induce vigorous microglial activation within 3 hours and lead to neuroinflammation for up to 10 months, despite that peripheral TNF-α levels already returned to baseline after 1 week. We also recorded robust microglial activation after 4 h as evidenced by morphological changes, upregulation of inflammatory gene expression, and activation of inflammatory pathways revealed by RNA-seq analysis. It remains to be determined why neuroinflammation persists after a single LPS challenge. It is plausible that epigenetic reprogramming leads to aberrant immunological imprinting in microglia (*42*).

Our transcriptome profiling from sorted microglia following the LPS challenge treated with or without TPT provides robust information regarding the mechanism of action of TPT in activated microglia. Firstly, among the top pathways from our GSEA analysis, several fundamental inflammatory pathways were significantly suppressed by TPT treatment, accompanied by the downregulation of many inflammatory genes. This further substantiated our data showing the effective anti-inflammatory property of TPT in inflammatory microglia as well as neuroinflammatory conditions in LPS and EAE models. Secondly, our GSEA analysis also revealed metabolic reprogramming in microglia following TPT treatment. We determined activation of pathways related to the mitochondrial respiratory chain, oxidative phosphorylation (OXPHOS), and steroid biosynthesis. Steroids have been shown to hinder the induction of pro-inflammatory cytokines in microglia and exert anti-inflammatory effects via ADIOL–ERβ–CtBP gene transrepression pathway (*43*). And it has also been well documented that a metabolic shift of microglia from glycolysis to OXPHOS with more oxygen consumption underlies the phenotypic change from a proinflammatory state to an immunoregulatory state, or a return to the homeostatic state (*44, 45*).

In addition, we also predicted potential transcription factors through which TPT could regulate downstream inflammation-related gene expression. Among these, *Ikzf1* and *Hdac2* are very intriguing. Interestingly, Ikaros (encoded by *Ikzf1*) regulates transcription by directly associating with HDAC-dependent complexes (*21*). As an epigenetic enzyme, the function of HDAC1and HDAC2 is redundant in adult microglia during the homeostatic state. An HDAC1/2 deficiency during development affects microglial survival, but they are not required for adult microglial survival. In contrast, its absence enhances the epigenetic modification of the DNA packaging protein Histone H3, as evidenced by increased H3K9ac and H3K27ac expression, and subsequently induces altered gene expression in microglia. This leads to a less pro-inflammatory milieu and enhanced amyloid phagocytosis (*46*). Furthermore, a hyperexpression of HDAC1/2 has been noted in several neuroinflammatory conditions, and inhibiting it has been proposed as a promising therapeutic strategy for controlling microglial activation (*47*). Ikaros was initially identified as a lymphoid-specific transcription factor, and its function in myeloid cells, especially in microglia, is under-appreciated (*48*). Several studies have examined the role of Ikaros in macrophages, but the results are inconsistent. In isolated tongue sole macrophages stimulated with LPS and lipoteichoic acid, Ikaros expression increased in a dose- and time-dependent fashion synchronized with increased expression of TNF-α and IL-1β, and was suggested to be a marker of inflammatory responses (*22*). The induction of Ikaros in LPS-stimulated macrophages and its fundamental role in activating NF-κB signalling was confirmed previously, and Ikaros thus regulates inflammatory responses (*49, 50*). However, Ikaros can also negatively regulate iNOS expression in macrophages, and loss of Ikaros may promote repolarization to proinflammatory macrophages (*51, 52*). Therefore, the role of Ikaros in macrophages and microglia is controversial and not fully understood. We identified Ikaros as a potential downstream target of TPT, but this needs more investigations in the future.

TPT treatment initiated at the peak of EAE failed to ameliorate clinical neurological signs of EAE. We consider that during the EAE recovery phase the infiltration of immune cells and progressive inflammation has retarded, while oligodendrocyte precursors proliferate at a high pace and differentiate into mature oligodendrocytes to repair demyelinated nerves. TPT treatment during this phase is not effective in this less inflammatory context but may affect cell proliferation and subsequent remyelination (*53*). Coincidentally, a very recent study using a network-based deep learning algorithm developed for novel drug target identification and *in silico* drug repurposing also predicted that TPT is an effective inhibitor of RORγt and thus, may cure EAE (*54*). In this study, they treated EAE mice from disease onset with TPT using a relatively high dose (10 mg/kg; i.p) and observed a protective effect. However, since TPT is a chemotherapeutic drug, with this high dose it is hard to distinguish if the effect is simply due to an overall immunosuppression/cytotoxicity on the highly proliferating immune cells. In our study, we used a much lower dose (1 mg/kg; i.p) and confirmed that our treatment regime did not induce cytotoxicity in naive mice. Furthermore, we also proved that inhibiting TOP1 only in CNS myeloid cells using TopoGami can improve EAE. At the same time, in addition to the *in vitro* validation of increased TOP1 expression in inflammatory microglia, we characterized an increased TOP1 expression in different neuroinflammatory contexts. To our knowledge, our study is the first to indicate an altered TOP1 expression in neuroinflammatory settings.

The *in vivo* application of classical compact, lattice-based DNA origami nanostructures has been hindered by two main challenges. Firstly, stabilization of this type of DNA origami nanostructures usually needs divalent cations, such as Mg^2+^, at the milli-molar range to overcome the electrostatic repulsion between closely packed DNA phosphate anions (*57*). This causes poor structural stability in physiological fluids which do not contain such a high concentration of cations. Secondly, bare DNA origami nanostructures would be rapidly cleared *in vivo* by nucleases (*58*). Thus, for reliable biomedical applications, the stability of DNA origami nanostructures under physiological conditions must be ensured. Our wireframe-stylized HR used in the work, due to its much lower DNA packaging density, has the advantage of cation independence. Recently, polymer protection strategies, such as oligolysine-PEG and cationic poly (2-dimethylamino-ethyl methacrylate, or PDMAEMA), have shown potential in stabilizing DNA origami nanostructures from nuclease degradation (*57, 59*). However, these polymers are not expected to possess any other biological functions. We now for the first time used β-glucan to coat DNA origami nanostructures. This represents a ‘two-birds-with-one-stone’ strategy achieving both the Dectin-1 receptor targeting on myeloid cells and the protection of DNA origami nanostructures to nucleases.

The construction and application of TopoGami enabled us to localize the effect of TPT on myeloid cells. Much to our surprise, from our *in vitro* experiments we observed that the nanostructures (HR and HR-Glu) as vectors themselves also decreased the inflammatory cytokine production in activated microglia to a certain degree, but the effect of TopoGami was far superior. We speculate that the nanostructure may have some sequences that act like NF-κB decoy oligos, which requires further investigation. Similarly, another study in which RAW264.7 macrophage cell line was treated with a tetrahedral DNA nanostructure also determined that although the DNA nanostructure induced mild cell activation towards an M1-like phenotype with increased iNOS, TNF-α, IL-1β, and IL-6 expression, it significantly attenuated the expression of these inflammatory cytokines in LPS/IFNγ-stimulated inflammatory macrophages through inhibiting LPS-induced phosphorylation of MAPK subfamilies (*60*). Unlike the effect *in vitro*, when we apply TopoGami and HR-Glu in the EAE model we only observed an improved EAE score in the TopoGami-treated group. We speculate that in the context of EAE, the CNS myeloid cells, unlike those homogeneously activated by LPS/IFNγ *in vitro*, are quite heterogeneous and may undergo variable activation states (*61*). The DNA nanostructure without TPT may attenuate the inflammatory response for some subsets while initiating the activation of other subsets. This may explain why HR-Glu only had some beneficial effects in cultured cells but not *in vivo*.

An unfulfilled challenge for future work is to modify TopoGami to pass through the blood-brain barrier, even though its integrity in patients with neuroinflammation is disrupted. Additional validation of TPT or TopoGami in other neuroinflammatory models such as the SOD1-G93A mice of ALS would further add support to our findings. Last, we briefly unveiled the potential involvement of Ikaros in microglia following TPT treatment, but the role of Ikaros in regulating microglial function warrants further investigation.

Our study indicates TOP1 inhibition as a promising therapeutic strategy for neuroinflammatory disorders. We not only further confirmed the important role of myeloid cells in neuroinflammatory contexts, but also provided insights and tools for developing appropriate future therapeutics, and thus may benefit patients with neuroinflammatory diseases, such as multiple sclerosis.

## Materials and Methods

### Study Design

The aim of this study is to determine the repurposing property of topotecan for the treatment of neuroinflammation with the application of DNA origami to achieve specific delivery to myeloid cells. Mouse primary bone-marrow-derived macrophages and primary microglia were used for in vitro experiments. The EAE and LPS model were induced for in vivo experiments; both adult males and females were tested in different repetitions (no gender difference was observed in terms of the effect of TPT or TopoGami). Mice were randomly assigned to each treatment group; the scoring of EAE mice, as well as the assessment of the explorative behavior of LPS mice, were performed in a blinded fashion. The sample size was determined by the availability of age-matched mice and previous experience with the EAE model and LPS model, and mice were excluded from the study under the following situations: 1) Mice had other problems and had to be sacrificed according to the veterinarian and the ethical permit, such as a long tooth, fighting and bleeding, rectal prolapse, infectious wound on the subcutaneous injection site of MOG. 2) In the EAE model, mice were dead or had to be euthanized before the start of any treatment. 3) For mice receiving the intracisternal injection, mice were excluded if they show signs of ataxia, imbalance, paralysis or death due to the unsuccessful injection. The *n* for the individual experiment was indicated in the figure legends, and the number of repetitions was also indicated. We defined and removed potential outliers using Graphpad Prism 8 with the built-in ROUT analysis (Q = 10%). Blinding was performed during data collection and analysis. Representative images from immunostaining were obtained from the subjects in randomly chosen areas.

### Ethics statement

Most animal experiments were performed at Karolinska Institutet, and are approved and performed in accordance with the Swedish National Board of Laboratory Animals and the European Community Council Directive (86/609/EEC) and the local ethics committee of Stockholm North under the ethical permits N138/14, followed by 9328-2019. The LPS-challenged mice survival experiment was performed at Fudan University where experimental protocols and animal handling procedures were approved by the Animal Care and Use Committee (ACUC) of Fudan University. For human brain tissue staining the rapid autopsy regimen of the Netherlands Brain Bank in Amsterdam (coordinator-I Huitinga) was used to acquire the samples with the approval of the Medical Ethical Committee of the Amsterdam UMC.

### Experimental animals

Male and female C57BL/6NTac mice (Taconic) were bred at the Comparative Medicine Department at Karolinska University Hospital, Sweden. Animals were maintained in a pathogen-free and climate-controlled environment with regulated 12 h light/dark cycles. All mice used for experiments were adults between 2 to 4 months of age, weighing 20–30 g, and had access to chow and water *ad libitum*.

### *Connectivity Map*-based drug screening

A public GEO dataset GSE76737 was utilized, which contains the gene expression profile of human microglia using a human gene ST 2.0 Microarray Chip (Affymetrix). The human microglia were polarized to either pro-inflammatory (M1) or immunoregulatory (M2a/M2c/Mtgfb) subtypes using the indicated stimuli: M1 (+ LPS/IFNγ), M2a (+ IL-4/IL-13), M2c (+ IL-4/IL-10/IL-13), and Mtgfb (+ TGFβ). The differential gene expression pattern from M0->M2a/M2c/Mtgfb and M1->M2a/M2c/Mtgfb were identified using the GEO2R tool. The top 1000 differential genes were selected; the up-regulated genes and the down-regulated genes were identified. Up-regulated and down-regulated genes from each polarizing pattern were converted to identifiers based on Affymetrix HG-U133A chip, followed by analysis using *Connectivity Map* (O2 version: broadinstitute.org/cmap). In each pattern, CMAP ranks the compounds using an algorithm based on their correlation to induce the pattern and gives a CMAP score for each compound with ‘+1’ indicating the strongest positive correlation and ‘-1’ being the strongest negative correlation. The GRP files needed for *Connectivity Map* input were provided in the Supplementary file.

### Primary cell culture

Primary bone marrow-derived macrophages and primary adult mouse microglial cell cultures were established based on our previous protocol with modifications (*62*). A detailed protocol is provided in the Supplementary materials and methods.

### Cell stimulation and treatment

To induce an inflammatory response in macrophages and microglia, we used 100 ng/mL LPS and 20 ng/mL IFNγ to incubate cells for indicated time depending on the experimental purpose; 1 μM camptothecin (CPT) or topotecan (TPT) were added to the cell culture 30 min before LPS/IFNγ stimulation. For flow cytometry analysis, cells were cultured in 24-well plates (5 × 10^5^ per well) for 24 h, and the supernatants were saved for cytokine analysis or nitric oxide assay; for western blotting, cells were cultured in 6-well plates (1 × 10^6^ per well) for 3/6/12 h; for RT-PCR, cells were cultured in 96-well plates (5 × 10^4^ per well) for indicated time points; for immunocytochemistry (fluorescence), cells were cultured in black 96-well plates (2 × 10^4^ per well) for 6 h.

### Cytotoxicity assay

The cytotoxicity of drugs on the cultured cells was measured by the release of LDH (an indicator of cytotoxicity) using a CytoTox 96 Non-Radioactive Cytotoxicity Assay (Promega, G1780) following the provided technical bulletin.

### Reverse transcription-polymerase chain reaction (RT-PCR)

Total RNA of cells or brain tissue (hippocampus and hypothalamus) were automatically isolated using QIAcube (Qiagen, 9001292) with the RNeasy mini kit (Qiagen, 74106) according to the product instructions and underwent 15 min on-column DNase digestion using an RNase-Free DNase Set (Qiagen, 79254). cDNA was prepared with reverse transcriptase using iScript kit (BioRad Laboratories, 1708891) following the standard protocol provided. Amplifications were conducted using SYBR green (BioRad Laboratories, 1708886) according to the manufacturer’s instructions and the 384-well plates were run in a BioRad CFX384 Touch Real-Time PCR Detection System. Primers were designed to work at approximately +60 °C and the specificity was assessed by melt curve analysis of each reaction indicating a single peak. The primers used for *Mus musculus* in this study are listed in Table S2.

### Nitric oxide detection and cytokine analysis

The measurement of nitric oxide (NO) in the supernatants was performed using the modified Griess reagent (Sigma, G4410). Samples (100 μL) were plated in a flat 96-well plate (Corning), and mixed with equal volumes of freshly prepared Griess reagent, and incubated for 15 min at room temperature. The plate was read at 540 nm using a microplate reader (LabSystems). The cytokine release to the supernatants was measured using BD™ Cytometric Bead Array (CBA) following the manufacturer’s protocol and detected using a BD FACSVerse flow cytometer. The following kits were used to detect indicated cytokines: Mouse IL-6 Flex Set (BD, 558301), Mouse TNF Flex Set (BD, 558299), Mouse MCP-1 Flex Set (BD, 558342), and Mouse IL-10 Flex Set (BD, 558300). The cytokine level in the supernatant was reflected by the median fluorescence intensity (MFI) of PE of the corresponding capture beads.

### LPS challenge

We followed a previous protocol and injected a single dose of systemic lipopolysaccharide (LPS, 5 mg/kg, i.p) which induces severe neuroinflammation (*13*). 10 μL of TPT (0.1 mg/kg) or PBS was delivered directly to the CNS before LPS injection via intracisternal injection under isoflurane inhalation. In another experimental setting, we also injected 10 ng LPS to the CNS via intracisternal injection and administered TPT (1mg/kg, i.p) to evaluate its effect on counteracting LPS-induced neuroinflammation. We also employed a lethal dose LPS challenge (18 mg/kg, i.p) based on a previous study to examine the overall protective effect of TPT on endotoxic shock (*63*). TPT (1mg/kg, i.p) was administered 1 h before a lethal dose of LPS challenge, and mice were kept for a maximum of 96 h.

### Explorative behavioral test

To reflect mice brain function following the LPS challenge, we evaluated novel object-evoked curiosity following the methods from previous studies with modifications (*64, 65*). Briefly, 24 h post-LPS challenge (5 mg/kg), mice were placed in an observation cage with a novel object (a fluffy teddy bear toy; diameter 5 cm) placed in the center of the cage; the behaviors were videotaped for 5 min. The cumulative number of times the mouse investigated the toy (e.g. touching, sniffing, trailing) was determined from the video records by a trained observer who was blind to the experiments.

### Induction of experimental autoimmune encephalomyelitis (EAE)

EAE was induced based on previous protocols in our lab with modifications (*66*). In brief, recombinant mouse myelin oligodendrocyte glycoprotein (MOG; amino acids 1–125 from the N terminus) was expressed in *Escherichia coli* (*E. coli*) and purified to homogeneity by using chelate chromatography following a previous protocol (*67*). The detailed protocol is provided in the Supplementary materials and methods.

### Intracisternal injection

We followed a previous protocol to perform intracisternal injection (*68*). In brief, mice were anesthetized under isoflurane inhalation, with the head bending forwards to expose the aperture between the occiput and the atlas. A 27G dental needle (Terumo, DN-2721) with the tip (around 3.5 mm) bent at an angle of approx 40° connected with a Hamilton syringe via a polyethylene tube was inserted into the cisterna magna. Up to 10 μL of the solution was slowly injected in approx 10 s.

### Flow cytometry

Single-cell suspensions were plated into 96-well V-bottom plates and stained at 4 °C for 20 min. Dead cells were removed using Live/Dead Fixable Near-IR or Yellow Dead Cell Stain Kit (Invitrogen, L34976 or L34959, 1:500) in each panel. Cells were acquired using a FACSVerse flow cytometer (BD Biosciences) or a Beckman Coulter Gallios Flow Cytometer and data were analyzed using Flowjo or Kaluza. The antibodies and staining protocols for the panels were described in the Supplementary materials and methods.

### Cell sorting

After myelin removal of brain homogenates, the cell suspensions were stained with the following antibodies: PE anti-mouse Ly-6C (BioLegend, 128008, clone HK1.4, 1:200), PerCP/Cyanine5.5 anti-mouse/human CD11b (BioLegend, 101228, clone M1/70, 1:100), APC anti-mouse F4/80 (BioLegend, 123116, clone BM8, 1:100), PE/Cyanine7 anti-mouse CD45 (BioLegend, 103114, clone 30-F11, 1:100), and Live/Dead Fixable Yellow Dead Cell Stain Kit (Invitrogen, L34959, 1.500). The CD11b^+^CD45^+^Ly6C^-^F4/80^int^ cells were sorted and collected using a SONY SH800 Cell Sorter. For magnetic-activated cell sorting of splenocytes, the spleens were mashed, followed by red blood cells removal using ACK lysis buffer (Gibco, A1049201). The cell suspension was first incubated with anti-CD11b MicroBeads (Miltenyi Biotec, 130-049-601), and the on-column CD11b^+^ cells were collected as the myeloid cells. The uncollected non-myeloid (CD11b^-^) cells were processed with the Pan T Cell Isolation Kit (Miltenyi Biotec, 130-095-130) following the manufacturer’s protocol, and the untouched T cells were collected, whereas the on-column cells were flushed and collected as the B cells.

### Next-generation RNA sequencing

Microglia were sorted directly into TRIzol reagent (Invitrogen, 15596-018) and kept at −80 °C before sending to BGI Genomics (Hong Kong, China) for RNA extraction and sequencing. The detailed protocol is provided in the Supplementary materials and methods.

### RNA-seq data processing

RNA-seq read quality was assessed using FastQC (https://www.bioinformatics.babraham.ac.uk/projects/fastqc/). Reads were trimmed using Trimmomatic (*69*) and mapped to mouse transcripts sequence (GRCm38), counted, normalized, and quantified (TPM, transcripts per million) by Salmon (*70*). The raw counting table was imported and summarized by tximport (*71*), followed by differential gene expression analysis with EdgeR (*72*). Significantly deregulated genes were identified by the false discovery rate lower than 0.05. Gene set enrichment analysis of gene ontology (GO) and Kyoto Encyclopedia of Genes and Genomes (KEGG) pathway was performed with ClusterProfiler (*73*). Transcription factor prediction was performed using BART (*74*) and HOMER (*75*) with the differentially expressed genes identified by EdgeR. All the scripts used for bioinformatics analyses are available at Github: https://github.com/KeyiG/LPS_TPT_RNA-seq.git. The RNA-seq data is available in the ArrayExpress repository, under accession numbers: E-MTAB-10679.

### Human brain tissue processing

Human brain tissues were acquired from MS patients diagnosed according to the McDonald criteria and age-matched controls without neurological conditions (*76*). Controls were identified and selected from a larger cohort based on their pathological and clinical profile. Some controls were excluded if they had medical records of neurological disorders, cancer, or other inflammatory diseases of the CNS. We followed the rapid autopsy regime of the Netherlands Brain Bank in Amsterdam (coordinator Dr. I. Huitinga) to obtain donor tissues. Participants or next of kin were informed and consented to the brain autopsy procedure and the use of their tissue for research purposes. Tissues were fixed in 4% paraformaldehyde, processed and paraffin-embedded. Tissues of MS brains were selected and classified based on the size and type of lesion for quantitative analyses. Identification of the lesions was acquired by immunohistochemistry for myelin proteolipid protein PLP to detect myelin loss and HLA-DR to detect myeloid cell activation (*77*).

### Gene Expression Omnibus data mining

Several datasets from Gene Expression Omnibus (GEO) were revisited and mined for examining TOP1 expression. GSE97930 and GSE73721 analyzed by the Brain Myeloid Landscape 2 (http://research-pub.gene.com/BrainMyeloidLandscape) were used for comparing TOP1 expression in different brain cell types in healthy humans (*78*). Datasets GSE85333 (from stimulated primary human macrophages), GSE108000 (from brain white matter of MS patients and healthy donors), and GSE18920 (from spinal cords of sporadic ALS patients or healthy donors) were analyzed with GEO2R, and the relative mRNA expression of *TOP1* was plotted.

### Statistical analysis

Statistical analyses were performed with Graphpad Prism 8, and a *P* value less than 0.05 was considered significant. In general, data were presented as group means ± s.e.m. The Student’s unpaired two-tailed *t*-test was used to compare the difference when two groups were compared, and one-way ANOVA with Dunnett’s Multiple Comparison test for comparison for more than two groups. For EAE score-based analyses (average, cumulative, and max EAE score), the Kruskal-Wallis test with Dunn’s multiple-comparisons test was used when more than 2 groups were compared, and the Mann–Whitney *U* test was used to compare between two groups. The area under the curve (AUC) of the EAE clinical course approaches continuous variable with normal distribution despite the EAE score being ordinal and was therefore compared using either one-way ANOVA with Dunnett’s Multiple Comparison test (more than two groups) or Student’s unpaired two-tailed *t*-test (between two groups). We specified the detailed statistical analysis for each graph in the corresponding figure legends.

## Acknowledgments

We thank Tojo James for converting the gene symbols to the Affymetrix HG-U133A probe identifiers. We thank Eliane Piket for introducing the techniques for intracisternal injection. We thank Sebastian Lewandowski for providing insights about ALS and related scientific discussions. We appreciate Feng Yang’s advice on the artwork design. We are grateful to the staff at our animal facility (AKM) for animal caretaking, and to people at our Neuroimmunology Unit, especially Mohsen Khademi, and Gunn Jönsson.

## Funding

StratNeuro funding for Collaborative Neuroscience Projects at Karolinska Institutet (RAH and BH)

Swedish Neurofonden Foundation (F2020-0017) (KZ)

China Scholarship Council 201700260280 (KZ) and 201700260271 (KG)

National Natural Science Foundation of China 82074538 (JW)

UPPMAX SNIC project 2020/16-223 and 2020/15-292 (CK)

## Author contributions

KZ conceived, designed, and planned this project; he also performed the *Connectivity Map* analysis as well as most of the experiments, with input from RAH, XMZ, AO, and HL; YW, KZ, RAH, and BH designed the β-glucan-coated DNA origami and TopoGami, and YW produced these nanomaterials and tested their basic properties under the supervision of BH. For RNA-seq, KZ sorted the cells and prepared the samples for sequencing; KG performed the RNA-seq data analyses, plotted the figures for publication, and uploaded the codes under the supervision of CK. HS performed the Western blotting experiments and helped with flow cytometry and sorting. EN performed the stainings on human brain tissues under the supervision of SA. The survival experiment following the LPS challenge was performed by JS under the supervision of JW; JS also tested TPT in a cuprizone-induced demyelinating model and JW also provided help with Connectivity Map. Master students SYK and CB learned and performed in vitro experiments under the practical supervision of KZ; SYK also helped with dissections, animal behavioral tests, and analyses, as well as scoring the EAE mice. JH induced MOG-EAE and scored the EAE mice. XMZ provided technical support on in vitro cell culture and analyses of flow cytometry data. AO produced MOG, performed experiments of tail blood analysis, and provided technical support for flow cytometry and in vivo experiments. KZ, YW, HS, and KG wrote the manuscript with suggestions from all authors, especially RAH for revising and editing the whole manuscript, and BH, AO, CK, HL for editing the corresponding part of the manuscript.

## Data and materials availability

All data associated with this study are present in the paper or the Supplementary Materials. RNA-seq results are available in ArrayExpress repository (E-MTAB-10679).

## Figures and Tables

**Extended Data Fig. 1.**
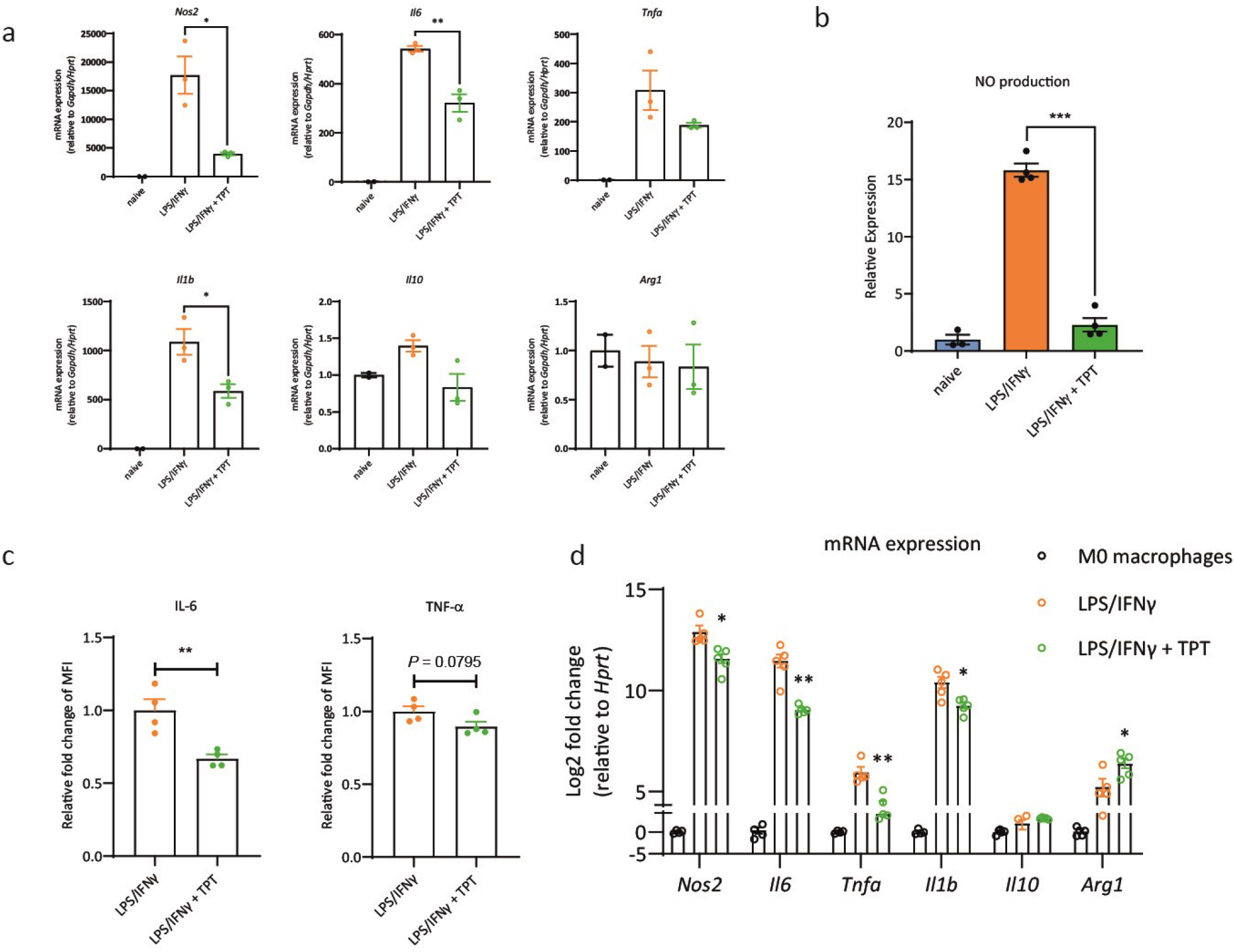
Topotecan (TPT) inhibits inflammatory responses in microglia and macrophages. **a**, mRNA expression of *Nos2, Il6, Tnfa, Il1b, Il10* and *Arg1* in microglia after 2 h LPS/IFNγ stimulation incubated with 1 μM TPT or control (PBS). *n* = 3 per group; naive resting microglia were included as a negative control. **b**, Nitric oxide release from microglia treated with 1 μm TPT or control under LPS/IFNγ stimulation after 24 h. *n* = 4 per group; naive resting microglia were included as a negative control. **c**, The cytokine release of IL-6 and TNF-α from stimulated microglia treated with 1 μm TPT or control was measured using cytometric beads assay; *n* = 4 per group. **d**, mRNA expression of *Nos2, Il6, Tnfa, Il1b, Il10* and *Arg1* in macrophages after 24 h LPS/IFNγ stimulation incubated with 1 μM TPT or control; *n* = 4-5 per group.

**Extended Data Fig. 2.**
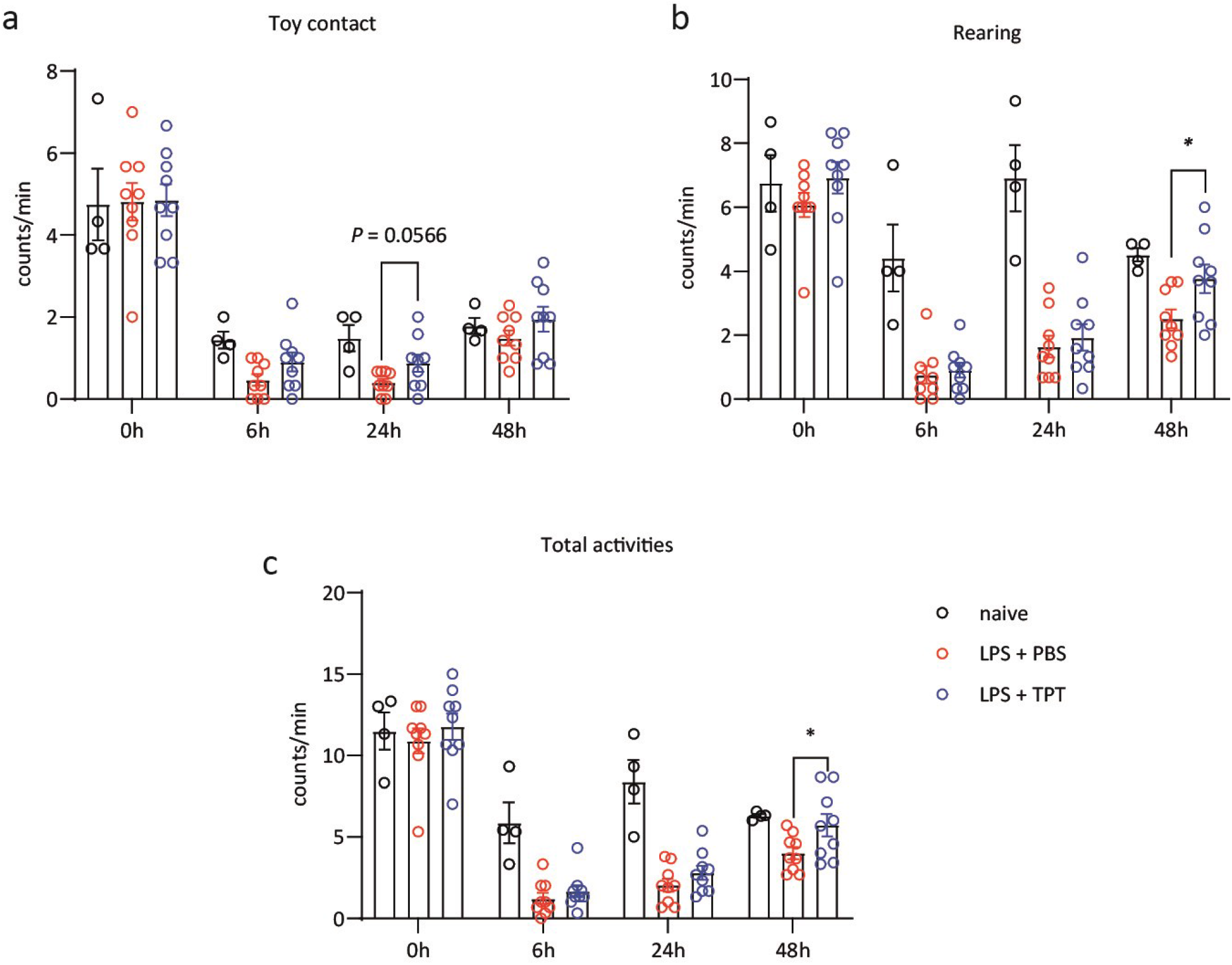
Cage explorative behavior of mice receiving LPS injection via intracisternal routine. 10 ng of LPS was injected directly into the CNS via intracisternal injection and 1 mg/kg TPT (or PBS) was administered intraperitoneally. The frequency of toy contacts (**a**), rearing (**b**) and total activities (toy contact + rearing) (**c**) was recorded during the following 48 h. *n* = 9 for PBS and TPT group, *n* = 4 for the naive control group. * *P* < 0.05. Statistical analyses comparing the LPS-PBS group with the LPS-TPT group were performed using Student’s unpaired two-tailed *t*-test.

**Extended Data Fig. 3.**
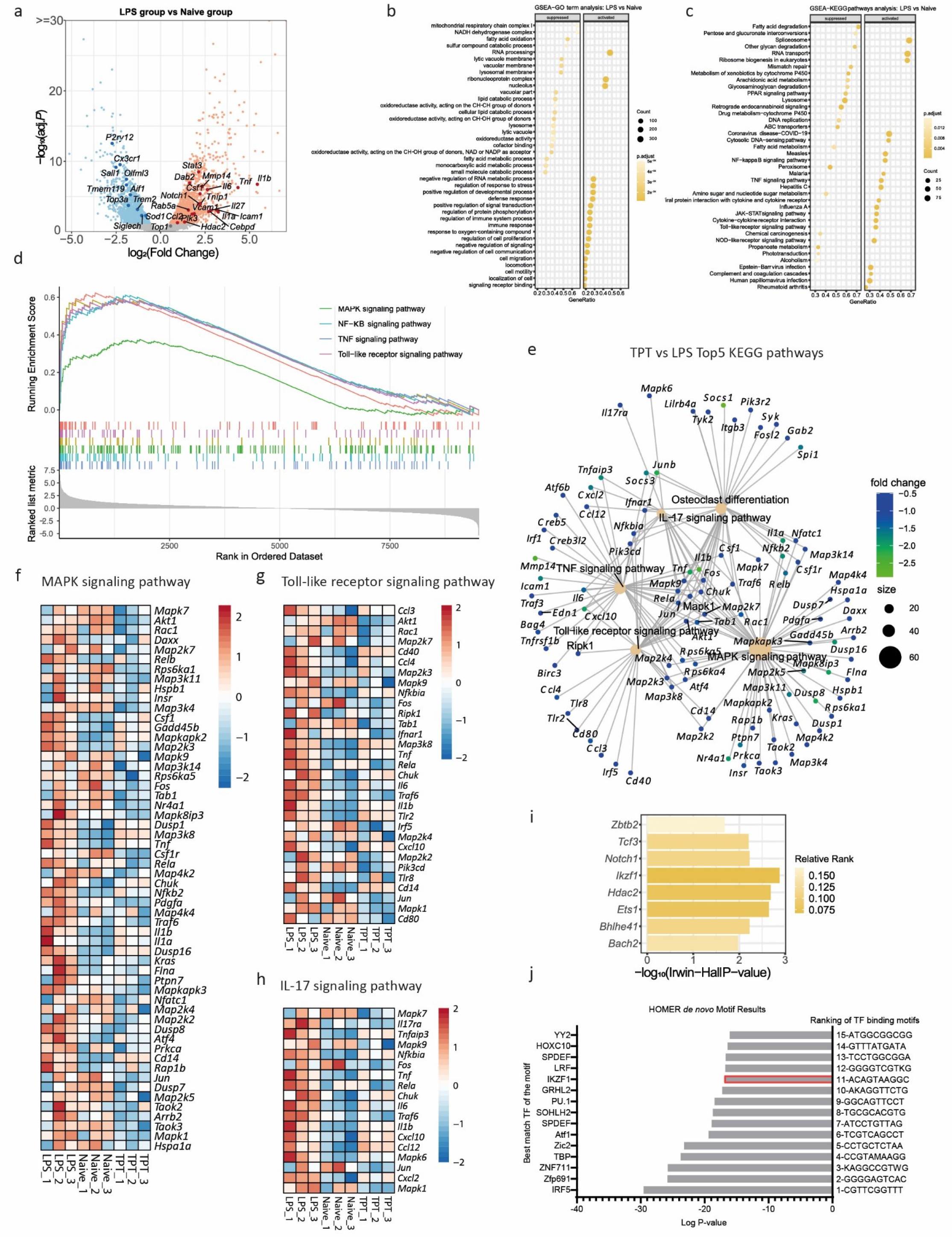
RNA-seq analysis of sorted microglia. **a**, Volcano plot depicting the DEGs between the LPS and naïve groups. Dots represent non-DEGs (grey) and DEGs (FC> 0, red; FC<0, blue); genes of interest were in dark colors. **b**,**c**, Bubble plots showing the top10 enriched suppressed/activated GO terms (**b**) as well as KEGG pathways (**c**) between the LPS and naïve groups. **d**, GSEA plot of the enriched KEGG pathways related to inflammation between the LPS and naïve groups. **e**, Category net plot of top5 enriched KEGG pathways when comparing the TPT group with the LPS group. The color gradient indicates the fold changes between the TPT and LPS groups computed by EdgeR (log2 transformed); the size of the category (pathways enriched) indicates the number of genes contributing to the enrichment. **f**-**h** Heatmaps plotting the changes in expression levels of the genes contributing to the enrichment analysis of MAPK signaling pathway (**f**), Toll-like receptor signaling pathway (**g**) and IL-17 signaling pathway (**h**) across samples when comparing the TPT group with the LPS group. **i**, Bar plot of the 8 genes in Fig. 4i according to BART result; Irwin-Hall *P*-value indicates the integrative rank significance; color gradients were coded by BART-computed relative rank. **j**, Bar plot of the *de novo* motifs enriched at the promoters of DEGs between the TPT and LPS groups; transcription factors with known motifs that matched the *de novo* motifs the best were listed accordingly.

**Extended Data Fig. 4.**
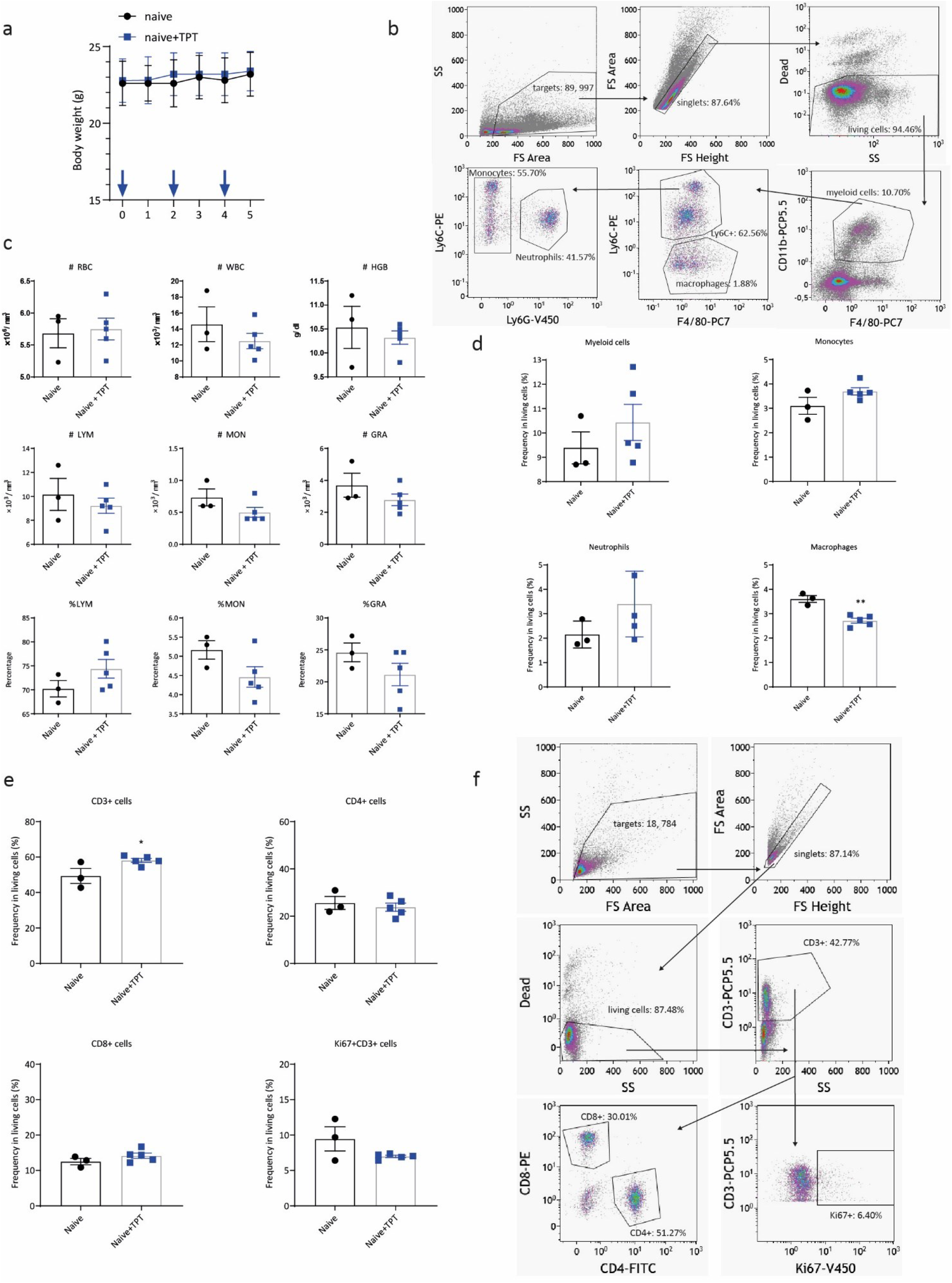
The effect of TPT treatment on the peripheral immune cell components under physiological conditions. **a**, The body weight change following 3 times TPT injection (every other day). **b**, On the following after 3 times TPT injection, the tail blood was sampled and analyzed using an ABC VET Animal Blood Counter. **c**, The gating strategy for the myeloid panel of the splenocytes. **d**, The frequency of myeloid components in the splenocytes 1 week after TPT treatment. **e**, The frequency of T lymphocytes in the splenocytes 1 week after TPT treatment. **f**, The gating strategy for the T lymphocytes.

**Extended Data Fig. 5.**
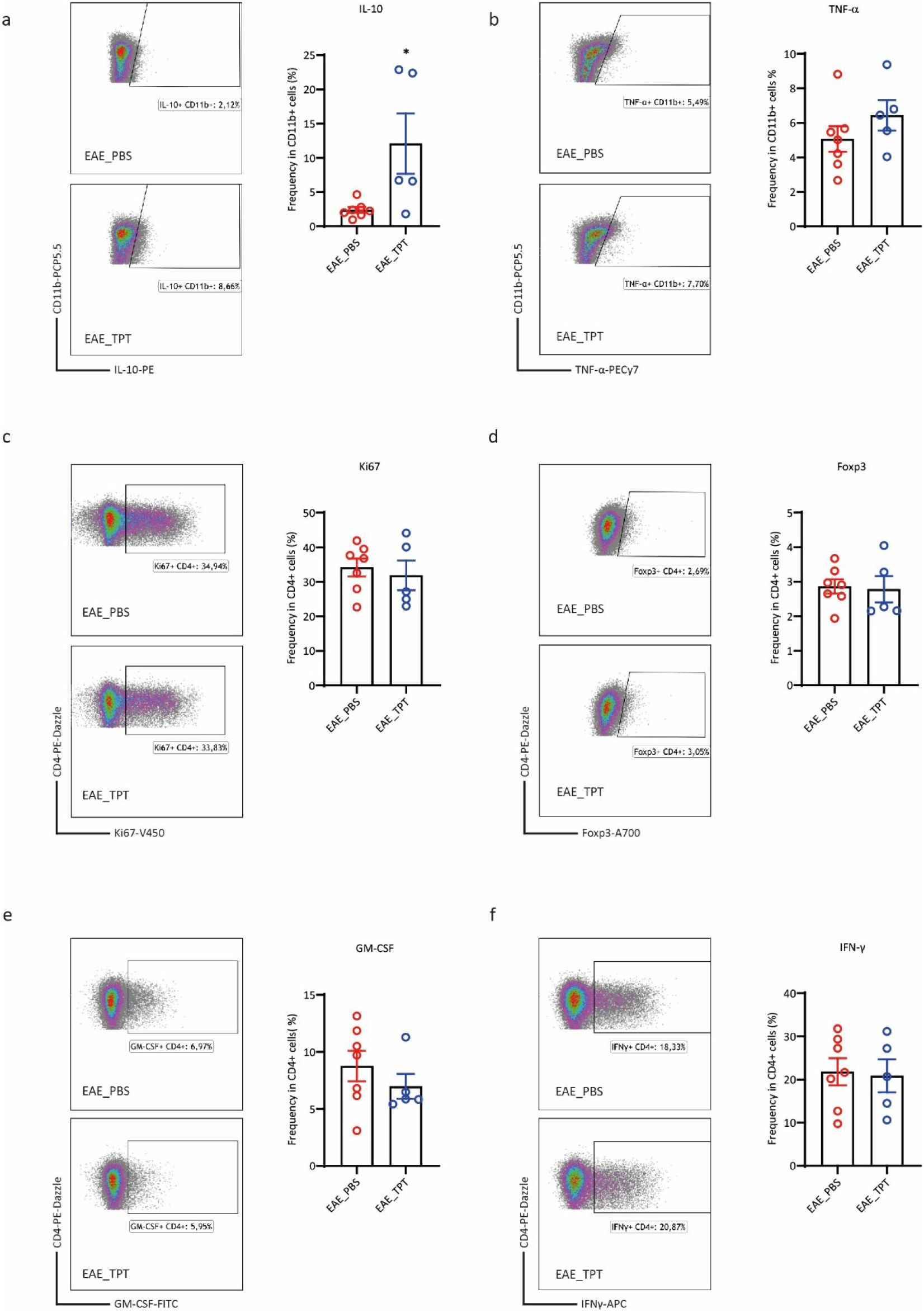
Cytokine production of immune cells from EAE spinal cords at disease peak. **a, b**, Flow cytometry plots showing the production of IL-10 (**a**) and TNF-α (**b**) in the CD11b+ myeloid cells; the frequency of IL-10-producing and TNF-α-producing CD11b+ cells were graphed. **c, d**, The frequency of Ki67+ CD4+ T cells (**c**) and Foxp3+ CD4+ T cells (**d**) was graphed. **e, f**, The production of GM-CSF (**e**) and IFN-γ (**b**) in the CD4+ T cells; the frequency of GM-CSF-producing and IFN-γ-producing CD11b+ cells were graphed. \**P* < 0.05 using Student’s unpaired two-tailed *t*-test; *n* = 5 and 7 for the TPT group and the PBS group, respectively.

**Extended Data Fig. 6.**
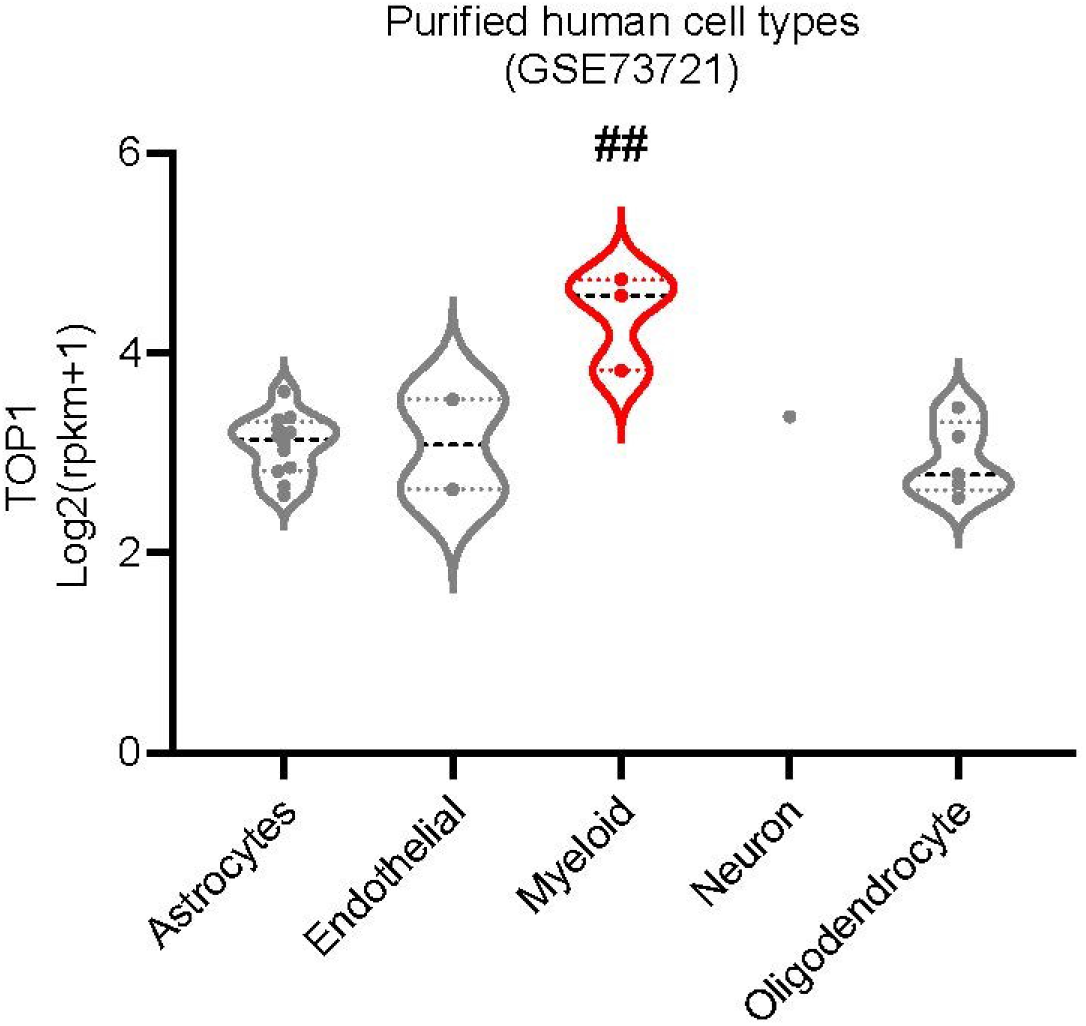
TOP1 mRNA expression in different brain cell types. Gene expression data from the dataset GSE97930 was extracted from the Brain Myeloid Landscape. ## *P*_*adj*_ ≤ 0.001 when comparing myeloid cells with other cell types, according to the analysis from the Brain Myeloid Landscape 2.

**Extended Data Fig. 7.**
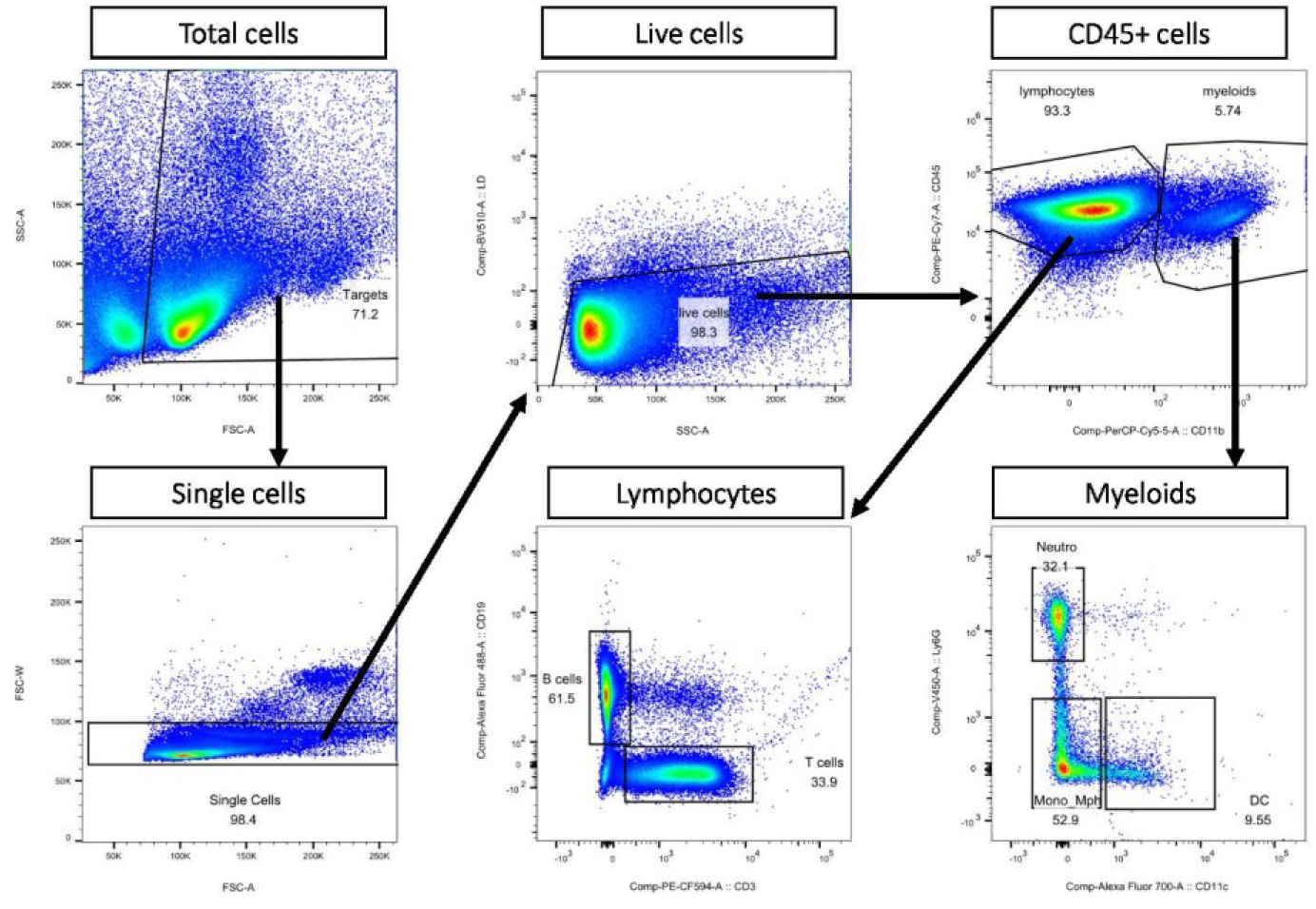
Flow cytometry gating strategy of the splenocytes.

**Extended Data Fig. 8.**
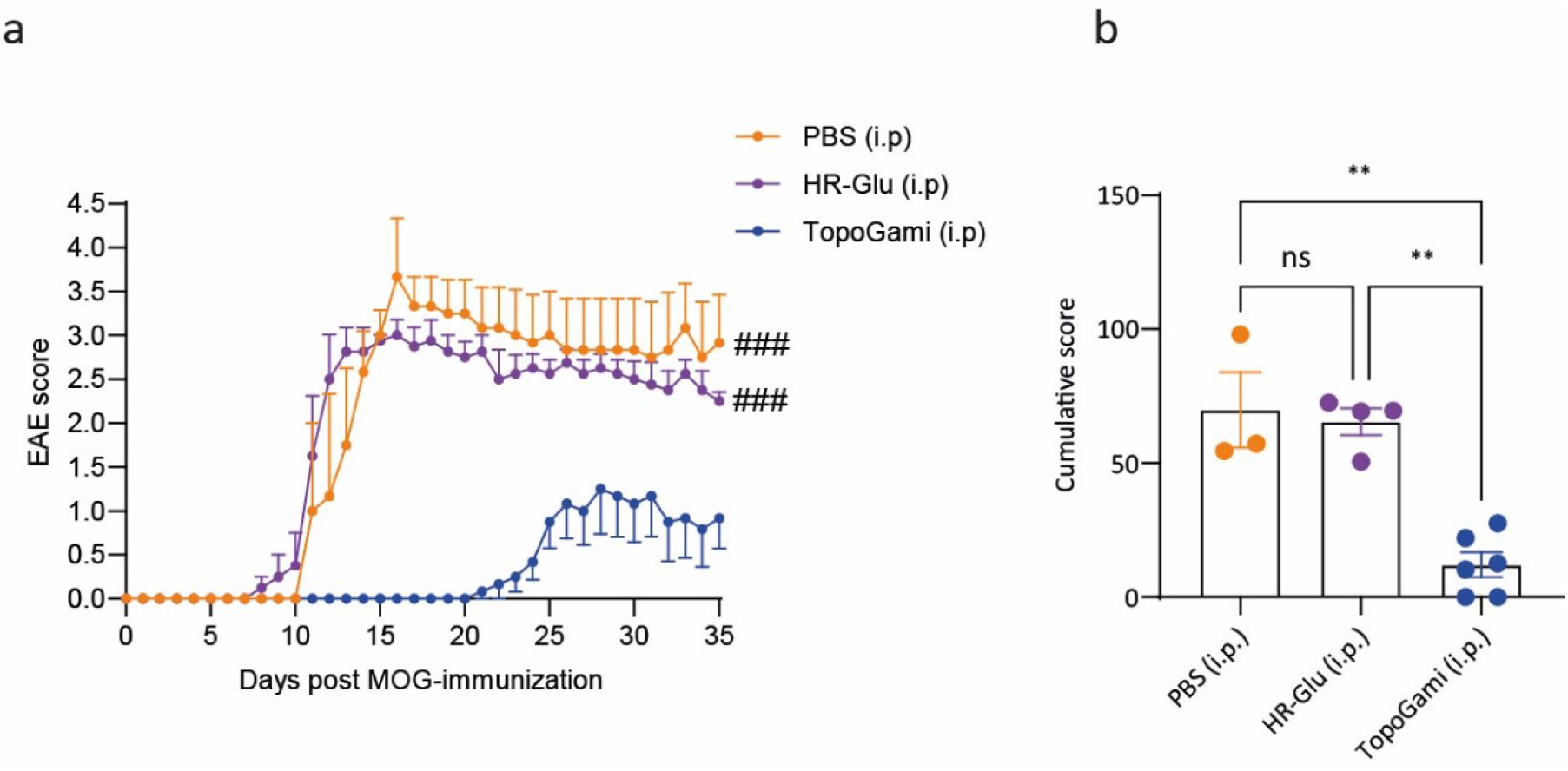
TopoGami treatment (i.p) prevents EAE progression. **a**, TopoGami (or HR-Glu/PBS controls) was administered to EAE mice at days 5, 7, and 9 post-immunization; the EAE clinical scores were graphed. **b**, Bar graph showing the cumulative EAE score in the 3 groups. ** *P* < 0.01 using Kruskal-Wallis nonparametric test for EAE cumulative score; ### *P* < 0.001 using one-way ANOVA with Tukey Multiple Comparisons Test for the area under the curve comparison of the EAE curves.

**Extended Data Table 1:**
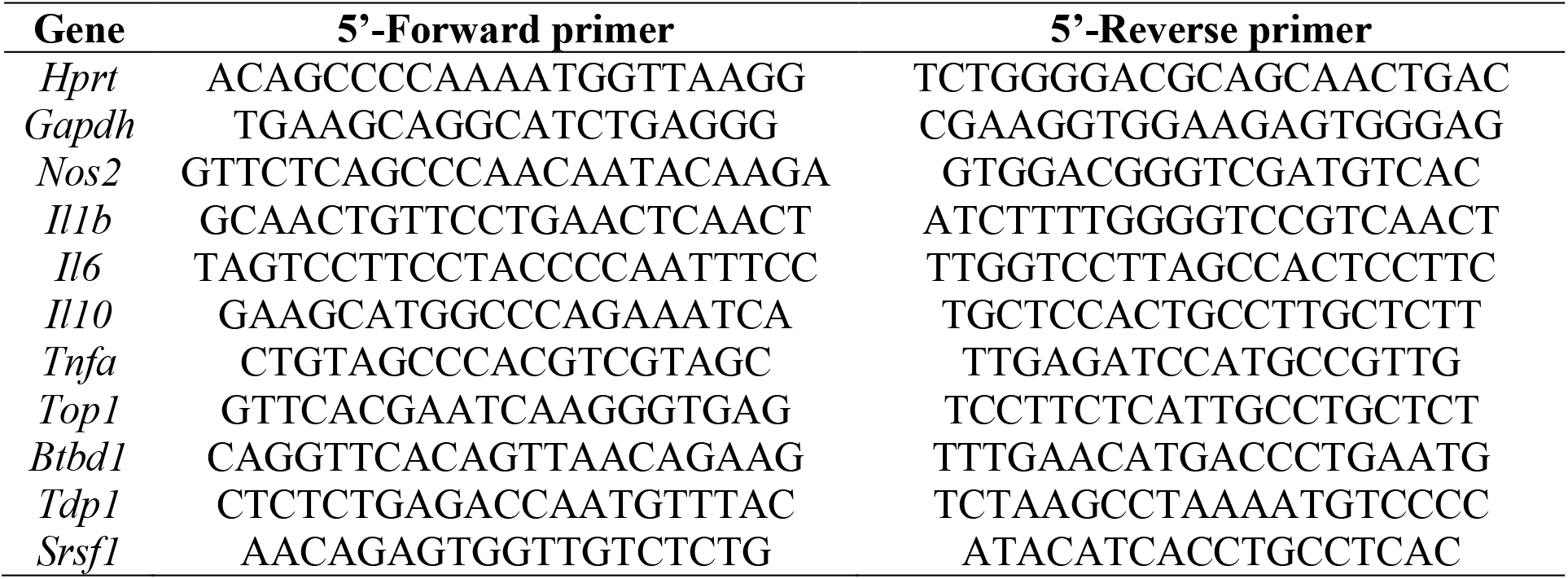
Primers used in the study (*Mus musculus*).

